# Multi-objective Engineering of Trimethylamine Monooxygenase for Improved Thermostability and Cofactor Use

**DOI:** 10.64898/2026.04.10.717641

**Authors:** Ruite Xiang, Martin Floor, Rasmus Ree, Albert Cañellas-Solé, Pål Puntervoll, Sergi Roda, Gro Elin Kjæreng Bjerga, Victor Guallar

## Abstract

Trimethylamine (TMA) is a major contributor to undesirable odours in protein hydrolysates derived from marine by-products, limiting their industrial use. Flavin-containing monooxygenases (FMOs) catalyse the conversion of TMA to the odourless trimethylamine N-oxide (TMAO); however, industrial applications demand enzymes that are both thermally stable and compatible with cost-effective cofactors. A thermostable variant of the *Methylophaga aminisulfidivorans* FMO (mFMO_20) can function at elevated temperatures but depends exclusively on the expensive and unstable cofactor NADPH. In this study, we investigated whether it is possible to simultaneously enhance thermostability and NADH compatibility using a multi-objective engineering strategy.

We first targeted residues in the cofactor binding site of mFMO_20 to restore NADH activity, which had been completely lost despite the wild type enzyme being naturally active with both cofactors. Variants derived from the thermostable scaffold partially recovered NADH activity but showed reduced NADPH activity. Given the wild type’s inherent NADH compatibility, we next pursued a stability-improvement approach, introducing highly conserved stabilizing mutations. This preserved cofactor competence but produced only modest improvements in thermostability. Finally, by combining physical, evolutionary, and statistical metrics, we obtained variants that retained higher NADPH activity after heat treatment than any previously reported thermostable mutants, while a subset also retained measurable NADH activity before heat treatment.

These findings show that combining complementary scoring strategies helps navigate the trade-off between stability and activity; while, robust NADH function under thermal stress remains elusive, with only one variant retaining detectable NADH activity after heat treatment, the results provide valuable insight into the underlying constraints linking stability and cofactor usage and highlights possible directions for engineering FMOs with both enhanced thermostability and cofactor compatibility.

**Author summary:** In this work, we aimed to improve an enzyme that could be useful in industrial applications but is limited by two common constraints: poor stability at high temperatures and dependence on an expensive cofactor. To make the enzyme more suitable for large-scale applications, we sought to engineer variants that are both more thermostable and compatible with a cheaper cofactor, NADH.

For enzyme engineering, we used a strategy that balances several properties rather than prioritizing a single trait. We combined tools that capture evolutionary patterns, protein physics, and AI-based predictions to explore which mutations might provide the right combination of stability and function. Through this approach, we obtained variants with improved heat resistance and higher cofactor activity retention.

## Introduction

Processing of marine by-products can yield highly nutritious products, but their use is often limited by an intense fishy odour caused by trimethylamine (TMA). This odour negatively affects consumer perception and hinders broader adoption in food applications [1, 2]. Enzymatic oxidation of TMA to the odorless trimethylamine N-oxide (TMAO) by bacterial flavin-containing monooxygenases (FMOs) represents a promising strategy, provided that the enzymes are sufficiently robust for industrial use [3]. Among these, the FMO from *Methylophaga aminisulfidivorans* (mFMO) is particularly attractive due to its high catalytic efficiency toward TMA [4]. Structurally, mFMO forms a homodimer, with each monomer composed of two domains: a larger FAD-binding domain and a smaller NADPH-binding domain. Despite its catalytic efficiency, the native enzyme exhibits only moderate thermostability and depends on the relatively costly cofactor NADPH, which limits its applicability under industrial conditions [5]. To improve thermostability, Goris *et al.* applied the Protein Repair One-Stop Shop (PROSS) algorithm [6] to mFMO, yielding several stabilized variants, with mFMO_20 emerging as the most thermostable. Despite this improvement, the continued reliance of mFMO_20 on NADPH remains a major bottleneck for large-scale implementation, even though a cofactor recycling system has been developed by coupling mFMO_20 with glucose dehydrogenase (GdhB) [7]. A key unresolved question is whether mFMO_20, or the native mFMO scaffold, can be engineered to utilize NADH as a more economical alternative cofactor, without compromising thermostability or catalytic performance.

Several computational strategies have been proposed for engineering cofactor specificity, including molecular dynamics-based approaches [8] structure-guided methods such as CSR-SALAD that identify positions for site-saturation mutagenesis [9], and machine learning-based models [10]. Despite these advances, cofactor engineering remains challenging, particularly when switching from NADPH to NADH. Indeed, approximately 89% of reported cases exhibit substantial losses in catalytic efficiency following cofactor switching [11], even when the mutations are introduced near the phosphate-binding site. This suggests that cofactor preference arises from the integrated architecture and dynamics of the binding pocket rather than from a single anchoring interaction.

Likewise, several physics-based tools exist which expedite screening for more thermally stable enzyme variants, such as PROSS [12], FRESCO [13] and FoldX [14] as well as machine learning tools such as ProteinMPNN [15]. While these approaches have proven effective, thermostability engineering can introduce mutations that compromise other enzyme functions, such as catalytic activity [16, 17] or cofactor affinity [18]. Consequently, achieving enhanced thermostability alongside altered cofactor preference requires balancing multiple, partially competing objectives, motivating the need for multi-objective optimisation.

In this work, we used a multi-objective optimization strategy to investigate whether NADH compatibility could be achieved in mFMO while improving thermostability. In doing so, we uncovered mechanistic evidence that this design problem is governed not only by static cofactor recognition, but by a tighter coupling between cofactor-binding orientation, reactive-state accessibility, and scaffold stability. Because similar trade-offs are likely to underlie cofactor use in other oxidoreductases, the framework developed here may prove broadly informative for enzyme cofactor-preference engineering.

## Results

### Evaluation of NADPH and NADH Activity in Wild Type and mFMO_20 Variants

It was recently shown that mFMO is able to use NADH as cofactor [19]. To determine whether the thermostable variant mFMO_20 retains this capability, we evaluated its catalytic activity using both NADPH and NADH as cofactors and compared the results with those obtained for the wild type mFMO. Consistent with prior reports, the wild type enzyme displayed robust activity with NADPH, achieving a turnover rate of approximately 82 µM product min^-^¹/µM enzyme or min^-^¹ under the tested conditions (Table 1). The wild type retained significant activity with NADH, albeit at a reduced rate (∼36 min^-^¹), representing roughly 44% of its NADPH-dependent activity. The mFMO_20 variant also retained substantial activity with NADPH, exhibiting a turnover rate of approximately 38 min^-1^, corresponding to about 45% of the wild type NADPH activity. This activity represents 73% of the reported by Goris et al. [5], which could be explained by differences in activity measurements and variation between expression batches. More markedly, when NADH was used as the cofactor, the activity of mFMO_20 dropped sharply to less than 1 µM min^-1^, representing only about 2% of the wild type NADH activity. These results indicate that while thermostabilisation using the PROSS design method preserved to some extent NADPH-dependent catalysis in mFMO_20, it severely compromised the enzyme’s ability to utilise NADH as an alternative cofactor. This pronounced loss of NADH activity led us to investigate the basis underlying cofactor binding and specificity in these enzyme variants.

### Molecular Simulations of Cofactor Binding in Wild Type and mFMO_20 Variants

To determine whether the mFMO_20 variant could be engineered to utilize NADH, the binding modes of NADPH and NADH in both mFMO and mFMO_20 were examined. PELE Monte Carlo simulations were conducted for mFMO and mFMO_20, each complexed with either NADPH or NADH. Preliminary PELE simulations (results not shown) indicated that the cofactors can adopt two distinct orientations within the binding pocket. Consequently, production simulations were initialised from both orientations to maximize sampling of each conformation and compare their relative stability. Analysis of the PELE landscapes demonstrated that both NADPH and NADH adopted multiple binding orientations (S1 Methods and S1 Fig). These orientations were categorized as NTA-oriented or ADE-oriented, depending on whether the nicotinamide or adenine moiety was facing the flavin (FAD) respectively, as defined by a C4N-N5 or C4A-N5 distance less than 5 Å, respectively (S1 Methods, Fig 1).

**Fig. 1.**
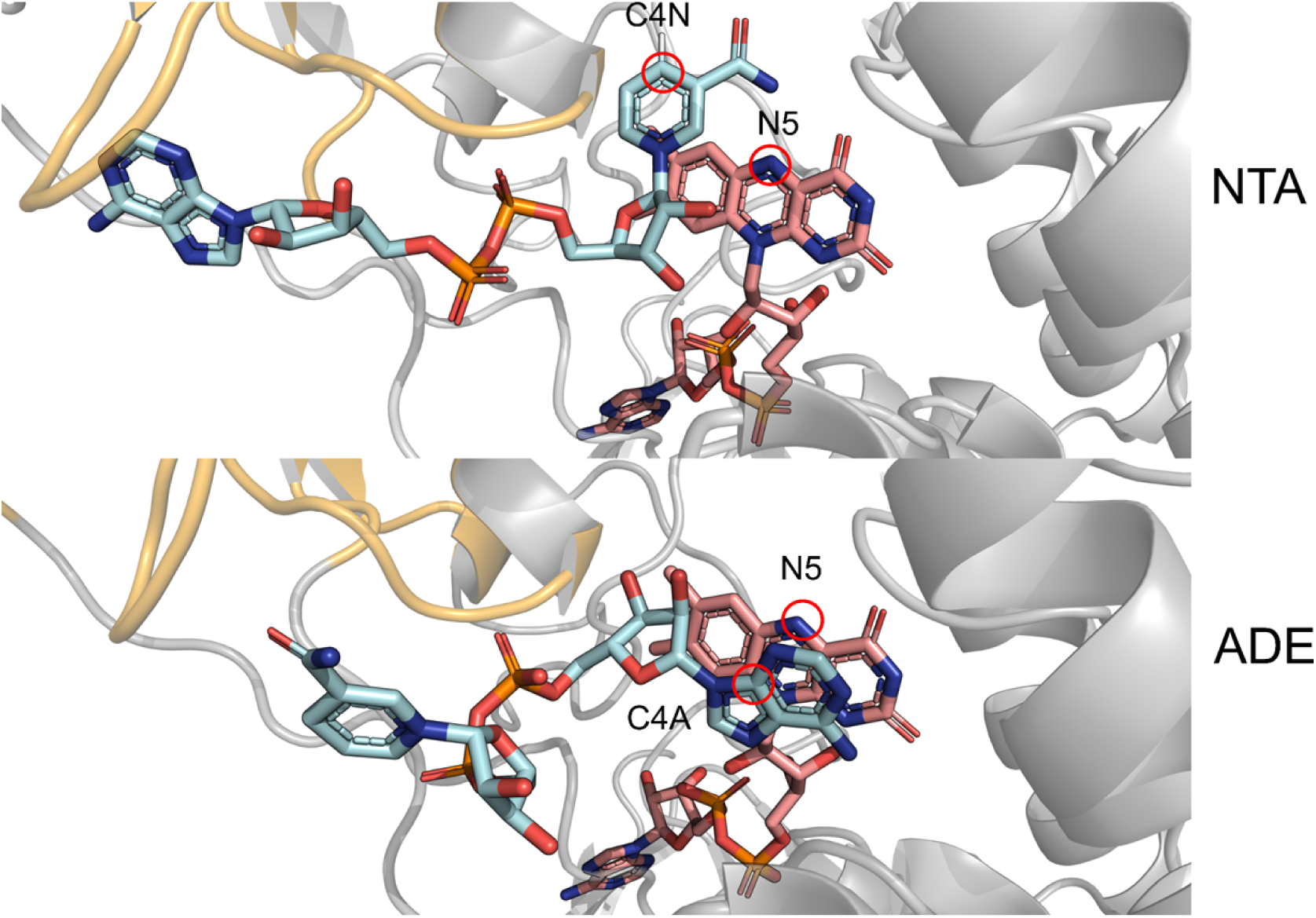
Representative NADH binding conformations in the active site of mFMO_20. Two possible binding orientations of NADH within the active site of mFMO_20 are shown. The large domain is depicted in gray and the small domain in yellow. FAD and NADH are represented as salmon and cyan coloured sticks, respectively. Nitrogen atoms are shown in blue, oxygen atoms in red, phosphorus atoms in orange, and carbon atoms in salmon or cyan according to the corresponding molecule. The atoms highlighted with red circles indicate the interatomic distances used to characterize the NTA and ADE orientations in the PELE and MD simulations.

To better visualize and interpret the simulation data, we derived free energy surfaces projected along these two nucleotide moieties distances (Figs. 2 and 3).

**Fig 2.**
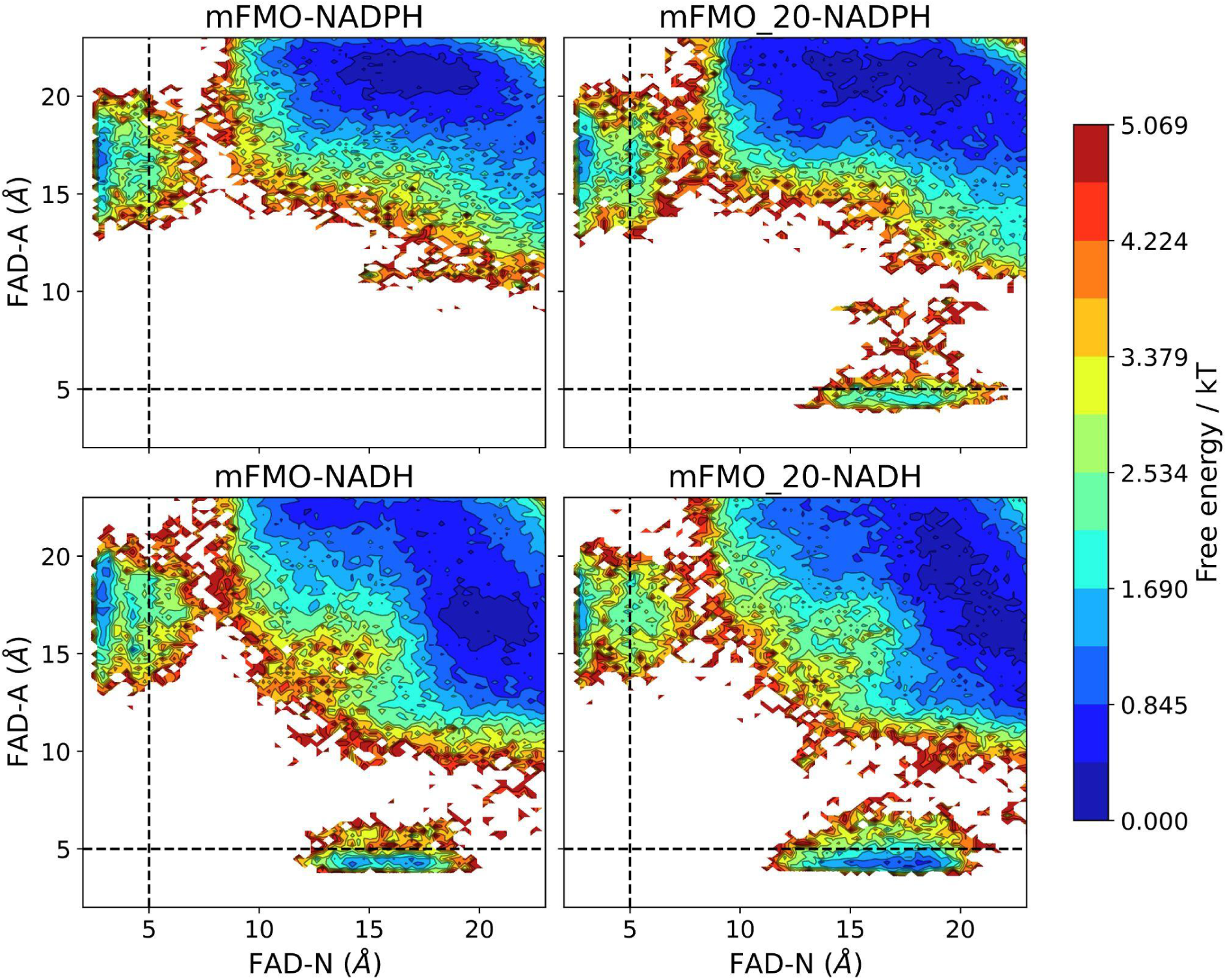
Free-energy surfaces of cofactor binding in wild type mFMO and the mFMO_20 variant. The free energy landscapes were obtained from PELE Monte Carlo simulations and are represented as a function of the distances between FAD-N (nicotinamide) and FAD-A (adenine) (A) mFMO with the NADPH cofactor, (B) mFMO_20 with NADPH, (C) mFMO with NADH, and (D) mFMO_20 with NADH. The black dashed lines indicate the distance threshold used to classify the orientations into NTA or ADE, 5 Å for both FAD-N and FAD-A.

**Fig 3.**
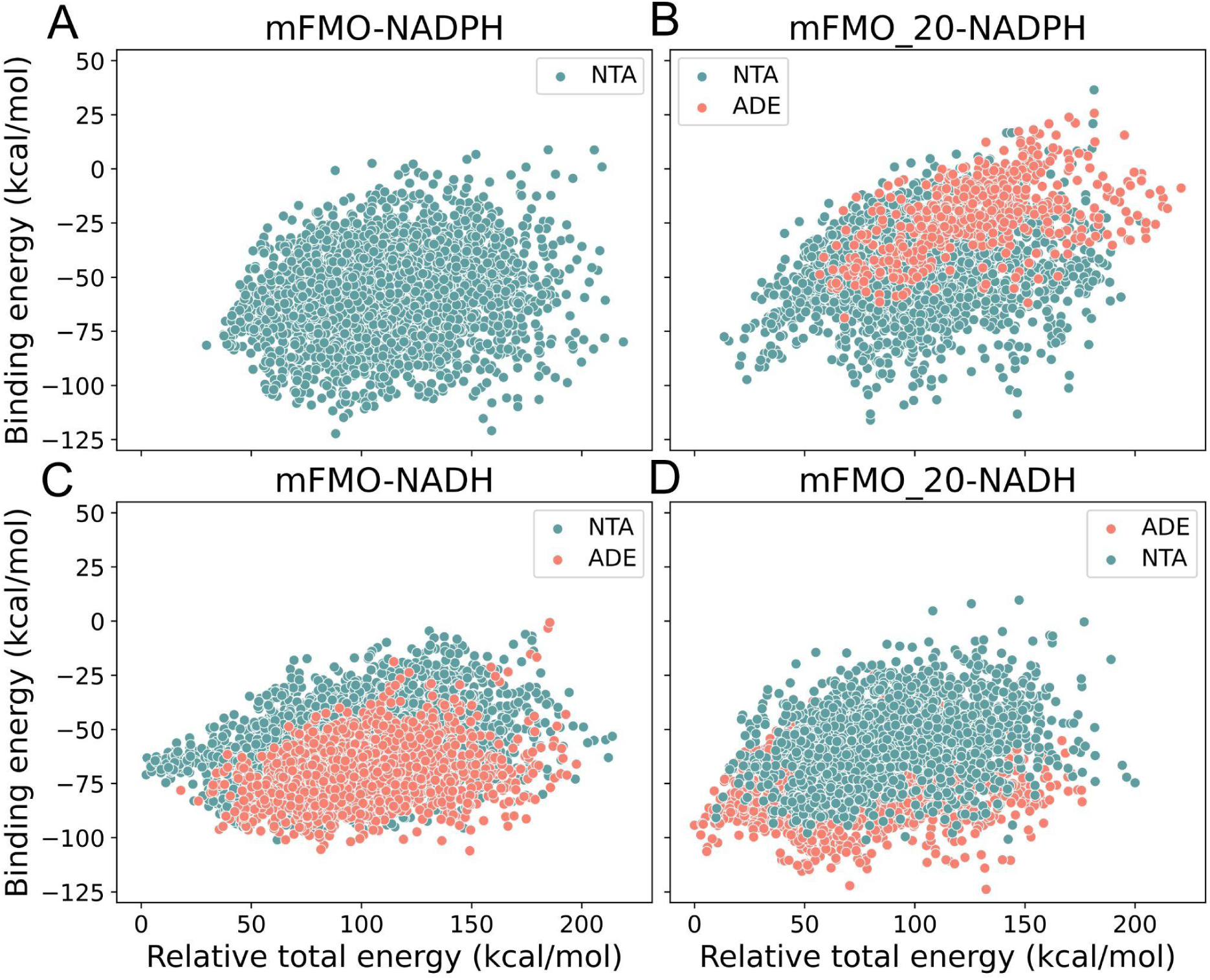
Scatter plots of binding energy vs relative total energy for mFMO and mFMO_20 in complex with NADPH or NADH. (A) mFMO + NADPH, (B) mFMO_20 + NADPH, (C) mFMO + NADH, and (D) mFMO_20 + NADH. The y-axis represents the binding energies (kcal/mol) for each conformation, while the x-axis represents the relative total energy, calculated as the total energy of each conformation minus the minimum total energy sampled. Conformations with FAD-N distances below 5 Å were classified as NTA, whereas conformations with FAD-A distances below 5 Å were classified as ADE.

In mFMO bound to NADPH, the free-energy surface (Fig. 2A) strongly favours NTA poses, with no ADE configurations observed. For mFMO_20 bound to NADPH, NTA poses show lower free energies than ADE orientations (Fig. 2B), consistent with the larger number of NTA conformations sampled (2.13% NTA vs 0.5% ADE poses). This preference is also reflected in the binding energy vs relative total energy analysis (Fig. 3B), where NTA poses show substantially more favourable total and binding energies than ADE poses, explaining why they are sampled more frequently.

In the presence of NADH, wild type mFMO shows a more favourable free-energy minimum for NTA orientations than for ADE (Fig. 2C), consistent with the larger number of sampled NTA conformations (2.55% NTA vs 1.06% ADE poses). This trend is also observed in the PELE total vs binding energy landscape, where the NTA configuration displays lower sampled total energies than the ADE configuration. However, both configurations show comparable minimum binding energies (Fig. 3C). In contrast, mFMO_20 bound to NADH (Figs. 2D and 3D) shows a preference for ADE orientations over NTA, with 1.80% ADE poses compared to 1.26% NTA poses. The ADE configuration also displays the most favourable binding and total energies among the two states, indicating a pronounced shift toward a non-productive binding mode in this thermostable variant.

Collectively, these results show an association between catalytic competence and the cofactor’s capacity to maintain a productive binding mode, with the nicotinamide ring positioned near the FAD isoalloxazine, in agreement with their experimental trend in cofactor usage. Both wild type mFMO and mFMO_20 predominantly adopt productive geometries with NADPH. The wild type scaffold is also capable of stabilising NADH in the NTA arrangement. In contrast, mFMO_20 preferentially stabilises NADH in the ADE-oriented mode, as consistently indicated by the free-energy landscapes and binding energies. These findings highlight the role of the NADPH 2′-phosphate in directing the binding ensemble toward a catalytically competent orientation. In the absence of this anchoring element (NADH), the systems explore a wider range of alternative orientations, including non-productive states, which can reduce the proportion of configurations suitable for catalysis.

### Multi-Objective Optimisation to Address Cofactor Binding Orientation

Given the shift in NADH binding orientation observed in mFMO_20, we explored whether restoring productive NADH binding through rational engineering would lead to an increased activity towards this cofactor. To this end, we applied a multi-objective optimisation (MOO) strategy, using a genetic algorithm to evolve enzyme variants with improved cofactor preference by targeting active-site residues near the cofactor (for details see S2 Methods).

Two optimisation objectives were established: (i) to maintain or improve overall structural stability, and (ii) to bias NADH binding toward the productive NTA orientation. The first objective was defined as the total Rosetta energy [20] of the NTA-oriented system, while the second objective was quantified as the binding energy gap between both orientations, calculated as the difference between the binding energy of the productive NTA pose and that of the non-productive ADE pose. This approach provides two objectives that reflect both stabilisation of the productive state and destabilisation of the flipped state, thereby indicating the relative preference for the catalytically competent orientation. These two objectives defined the Pareto front used by the genetic algorithm to prioritise variants that favour productive NADH binding while preserving structural integrity.

Following 100 genetic-algorithm iterations, the MOO generated variants exhibiting greater energetic separation between the productive (NTA-oriented) and non-productive (ADE-oriented) NADH states (Fig. 4). Across successive generations, the Pareto front advanced toward solutions that stabilized the productive orientation, as indicated by reduced energies for the NTA conformation and larger total and binding energies for the ADE conformation. This trend aligns with an optimisation trajectory that prioritizes stronger, catalytically relevant NADH interactions while maintaining the stability constraint.

**Fig 4.**
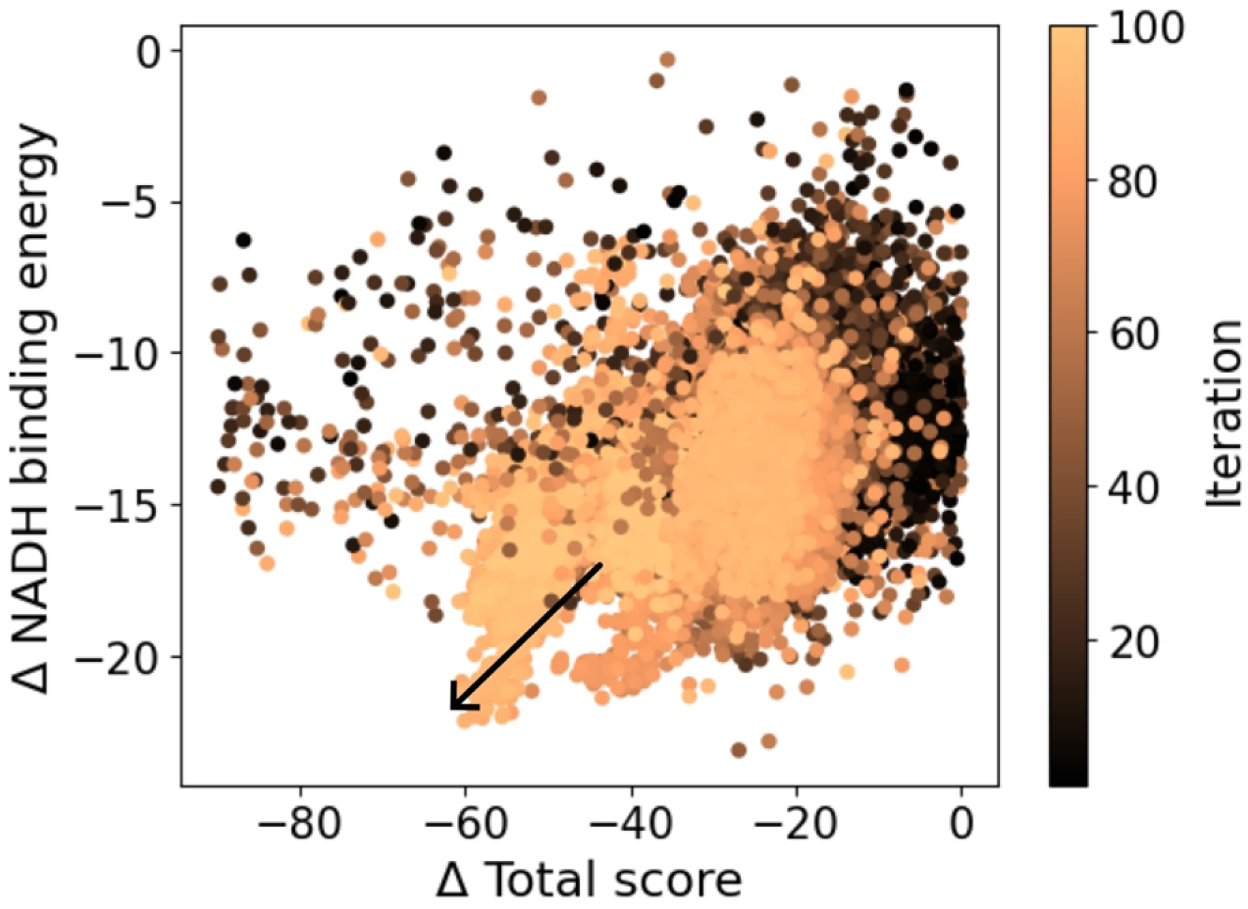
The results of the MOO run with mFMO_20 bound to NADH after 100 iterations. The arrow indicates the direction of the evolution towards better relative binding energies (y-axis) and total energies (x-axis) of the active conformation (NTA) of NADH compared to the inactive (ADE) one. The energy differences increased as the number of iterations (colour bar) of the genetic algorithm’s generations increased.

This MOO strategy was compared with AsiteDesign, a Rosetta-based protocol from our group that focuses exclusively on stabilising the productive NTA conformation without explicitly considering the flipped pose [21]. Additionally, a separate, manually guided approach was implemented by introducing mutations at residues adjacent to the phosphate-binding region, which are frequently targeted in cofactor-specificity engineering studies [9] (S2 Methods).

To prioritise variants for experimental testing, PELE was employed to evaluate how each design altered the NADH orientation landscape. For each design, both the productive NTA-oriented and non-productive ADE-oriented configurations were explicitly sampled, and their binding energies were compared to quantify the energetic preference between states. Variants exhibiting the strongest energetic bias toward the NTA orientation were selected as candidates, based on the rationale that stabilising the productive registry or destabilising the flipped registry should increase the proportion of catalytically competent binding configurations and thereby enhance NADH-supported activity (S2 Fig).

Twelve variants were selected for initial experimental testing: one manually guided design (BSC001), seven AsiteDesign variants (BSC002-BSC008), and four from the MOO run (BSC009-BSC012), with mFMO and mFMO_20 as controls. Activity assays demonstrated that most designs exhibited negligible NADH turnover (S3 Fig). Three variants, BSC009 and BSC012 (MOO) and BSC006 (AsiteDesign), showed measurable activity, surpassing the low residual NADH activity of mFMO_20. In contrast, none of the tested variants maintained substantial NADPH activity (S1 Table), suggesting that the design interventions introduced mutations that introduced a tradeoff that eliminated or significantly reduced NADPH-supported turnover.

### Optimisation for Thermostability and NADH Compatibility

Given that wild type mFMO exhibits measurable NADH activity, whereas mFMO_20-derived designs largely failed to recover it, we investigated whether thermostability could instead be introduced into the native scaffold while preserving its intrinsic cofactor-usage plasticity. Because the initial designs explored a broad, unconstrained mutational space and coincided with a marked loss of NADPH-supported turnover, we adopted a more conservative strategy that restricted the search space to minimise disruption of cofactor recognition and active-site geometry. In parallel, we monitored interaction energies with FAD, NADH, and the dimer interface (second subunit) throughout the evolutionary trajectory, enforcing near-wild-type values to preserve key contacts while accumulating stabilising substitutions.

This approach allowed us to test whether thermostability gains could be achieved without compromising cofactor-dependent activity, while enabling a direct comparison to previously reported PROSS-designed mFMO variants [5], which delivered substantial thermostability improvements. To implement this strategy, we constructed a mutant library using a PROSS-like workflow (S3 Methods). Given the mechanistic complexity of this enzyme, where productive function depends on tightly coordinated interactions between FAD and NAD(P)H across the flavin-reduction, oxygen-activation, and cofactor-recycling steps, and motivated by the observation that mFMO_20 improved stability yet lost NADH activity, we enforced stricter conservation filters than in standard stability design. Specifically, we restricted substitutions to those most frequently observed among natural homologs, with the aim of minimising perturbations to essential cofactor-binding and catalytic interactions. Subsequently, we conducted the optimisation in 10 independent runs of 100 iterations each, designating Rosetta total energy as the primary optimisation objective and evaluating NADH binding energies to remain close to wild type values as a post-optimisation filter (Fig. 5).

**Fig 5.**
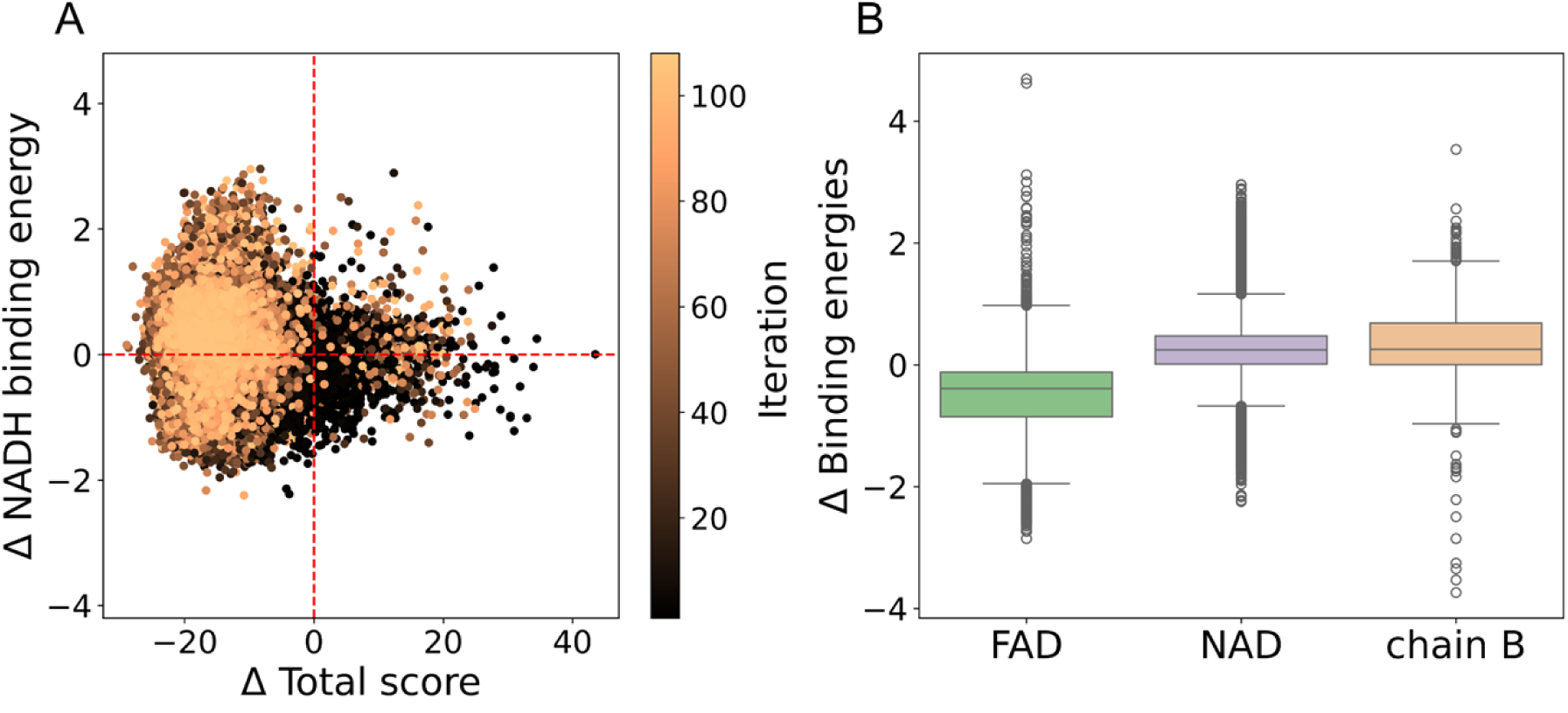
Optimisation of wild type mFMO for thermostability and NADH compatibility. (A) Scatter plot of ΔRosetta total score versus ΔNADH binding energy across ten independent genetic algorithm runs (100 iterations each) relatoce to the WT values. Each point represents a variant, coloured by iteration number from early (dark) to late (light). Red dashed lines mark the wild type reference (Δ=0). Variants in the lower-left quadrant show improved total stability and maintained or improved NADH binding. (B) Boxplots of interaction energy changes (Δ binding energies) for FAD, NADH, and the second enzymatic subunit (chain B) relative to the wild type. The inclusion of cofactors and the dimer interface enabled monitoring of stability across the full enzymatic assembly.

Ten representative designs were selected from this run based on a balance of low Rosetta energy, sequence diversity, and favourable stability signatures from molecular dynamics (MD) simulations, including RMSD, RMSF, and preservation of native contacts, as computed for the monomer and FAD constructs (S3 Methods). Experimental testing demonstrated that three variants (BSC018-BSC020) exhibited higher NADH and NADPH activities than mFMO_20 prior to heat treatment. However, following heat treatment, activity decreased for most designs; only BSC020 retained detectable NADPH turnover, although this activity remained below the post-heating activity of mFMO_20 (Fig. 6 and S2 Table).

**Fig 6.**
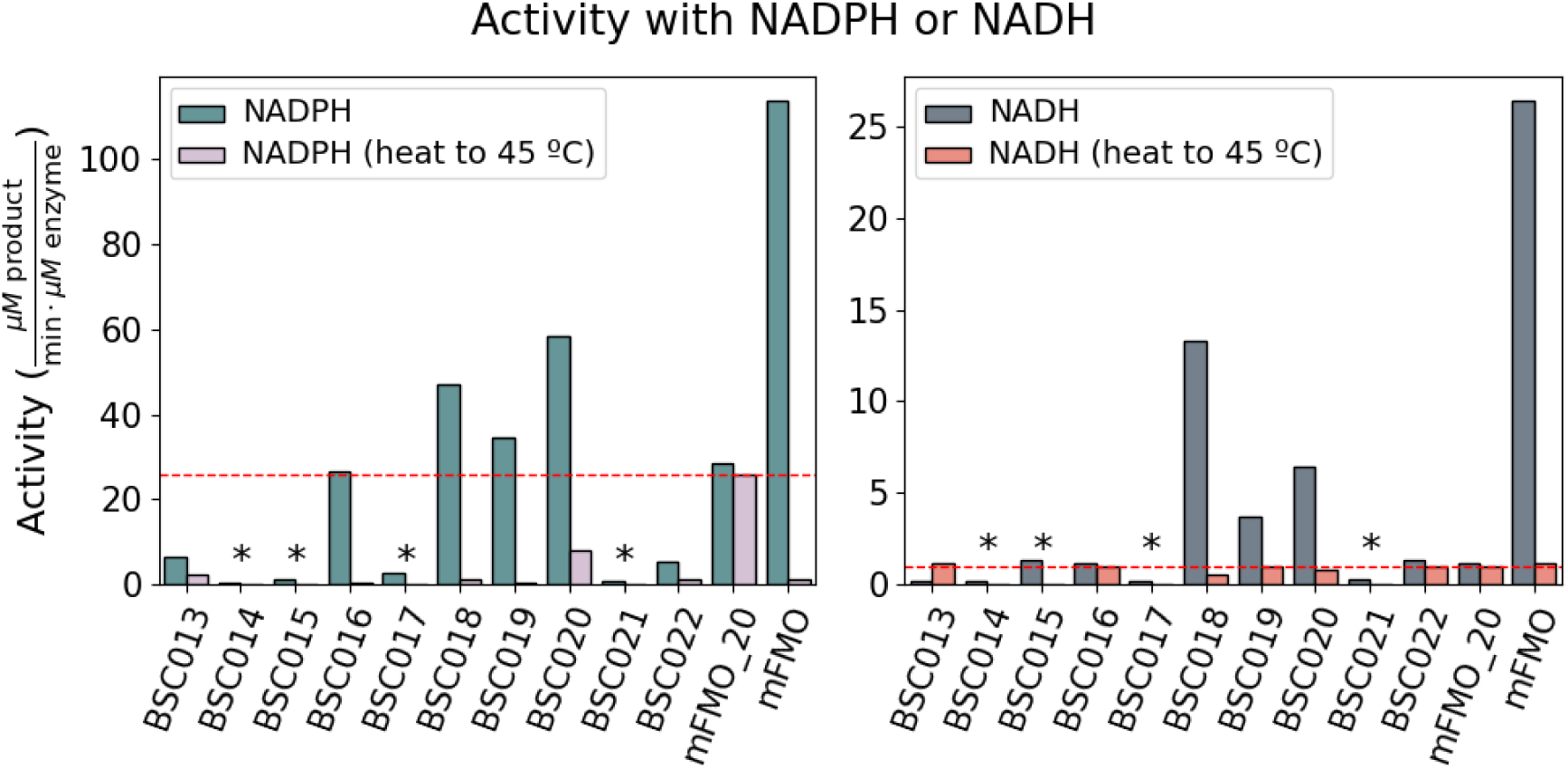
Activity of the mFMO variants from the second optimisation. Wild type mFMO and the indicated mFMO variants (50 nM) were purified, and enzyme activity was calculated using the indicated cofactor (either NADPH or NADH at 100 µM) and TMA (100 µM), monitoring the reaction at 340 nm (n=2) over 1 minute, either after heating at 45 °C or with no heat treatment. Variants 14, 15, 17, and 21, marked with asterisks, were not tested after heating due to low activity. The red dashed lines represent mFMO_20’s activity after heating.

Sequence analysis showed that substitutions across the ten selected variants were predominantly hydrophobic (61%), reflecting the compositional bias of the conservation-restricted library. In contrast, the PROSS mutant library used to generate the mFMO variants (S1 Appendix) exhibits a lower hydrophobic content (37.7%), favouring polar and charged substitutions capable of forming stabilising hydrogen-bond and salt-bridge networks. This broader interaction repertoire likely contributed to the substantial thermostability gains observed in mFMO_20. Notably, despite Rosetta total energies suggesting that our variants were, on average, more stable (S4 Fig), these scores did not correlate with improved thermostability, indicating that they do not fully capture its underlying determinants.

Accordingly, while our conservation-restricted strategy better preserved activity, it resulted in only modest stability gains, highlighting how the definition of the mutational landscape shapes the stability-function trade-off. Achieving larger stability gains may require expanding the accessible mutational set while retaining those combinations that preserve the underlying dynamical architecture supporting cofactor binding and turnover, an aspect that is unlikely to be captured by single-site frequencies alone.

### Multi-Objective Optimisation with Evolutionary and Language Model Scores

To overcome the limited stability gains from conservative mutations alone, we expanded the design framework beyond Rosetta total energies, incorporating additional evolutionary and statistical metrics and adopting the less restricted PROSS mutational library space. Potts model energies were used to capture residue co-variation patterns from natural homologs, reflecting both structural and functional couplings [22]. On the other hand, the ESM language model [23] scores (ΔLL or loglikelihoods and ΔP or sum of per-residue probabilities, see methods) were applied during ranking to estimate mutational fitness from large-scale protein data. We also incorporated structural features relevant to thermostability [24], such as the number of salt bridges and the solvent-accessible surface area (SASA) of exposed hydrophobic residues.

We applied this protocol in 10 independent replicas of 100 iterations for this third design campaign. Relative to the second Rosetta-only run, reopening the mutational space with a more hydrophilic library and co-optimising Potts and Rosetta energies redirected the evolutionary trajectories toward sequences with lower Potts energies (Fig 7), consistent with increased evolutionary compatibility, while also achieving lower Rosetta total scores (S5 Fig). Notably, although ESM scores were not included as explicit objectives, they improved in parallel with Potts energies, suggesting that the two sequence-based metrics capture partially overlapping, yet complementary, constraints on mutational plausibility (Fig. 7).

**Fig 7.**
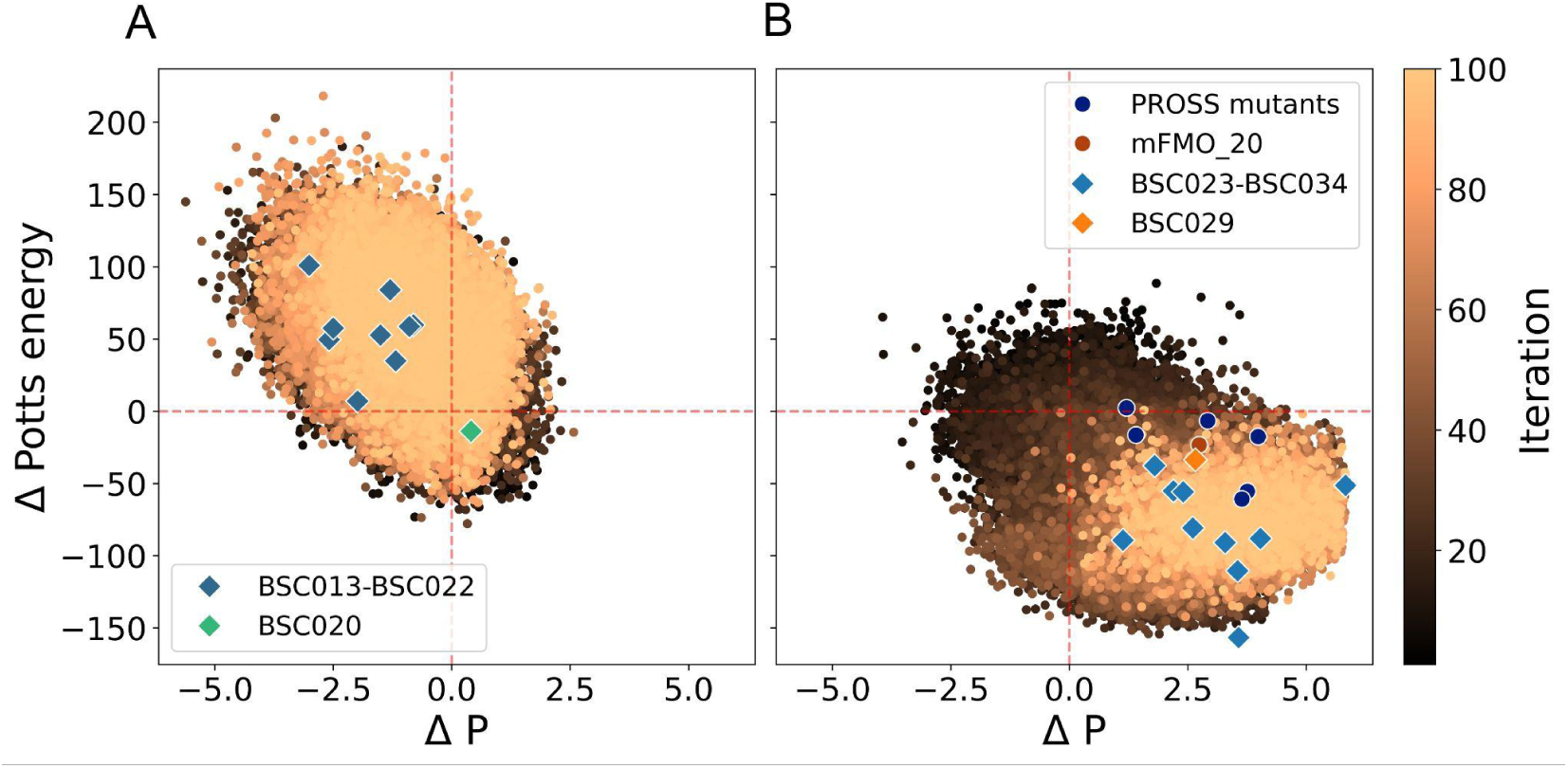
Results from the second and third optimisation campaigns. (A) Results from the second Rosetta-only optimisation run overlaid with the BSC013-BSC022 mutants; BSC020 is highlighted separately. (B) Results from the third optimisation run overlaid with the PROSS mutants (sequences obtained from [5] and analysed following the same scoring metrics) and the BSC023-BSC034 mutants; BSC029 and mFMO_20 are highlighted separately. The y-axis shows Δ Potts energies relative to the wild type, and the x-axis shows the Δ sum of probabilities or P. The colour bar indicates the number of genetic algorithm generations. Red dashed lines denote the wild type reference (Δ = 0).

From this engineering campaign, we selected twelve variants (BSC23-BSC034) for experimental validation, prioritising low Potts energies, favourable ESM scores, and overall sequence diversity (S4 Methods). As shown in Fig. 7, these twelve mutants span a range of favourable Potts energies and ESM scores. Notably, variants from the second optimisation run that did not exhibit improvements in thermostability (BSC013-BSC022, except BSC020) tend to display unfavourable Potts energies and ESM scores, while the PROSS variants reported in [5], all of which are thermostable, consistently show favourable scores. Together, these observations suggest that Potts and ESM scores are informative for prioritising variants within a thermostability-compatible region of sequence space.

Upon heating at 45°C, wild type mFMO lost all detectable activity, whereas most designs retained substantial NADPH turnover (Fig. 8 and S3 Table). Notably, four variants (BSC023, BSC026, BSC029, BSC031) exceeded mFMO_20 in post-heating NADPH activity, indicating improved activity retention under thermal challenge. Among these, BSC029 was the strongest performer: it retained 64% of the wild type NADPH activity after heat treatment, compared with 20% for mFMO_20 (S3 Table). Accordingly, BSC029 displayed only a ∼2% activity decrease upon heating, whereas mFMO_20 decreased by ∼26%, consistent with markedly enhanced thermal resistance. Four variants (BSC025, BSC026, BSC028, and BSC032) exhibited measurable NADH activity prior to heating, and notably, BSC025 uniquely retained detectable NADH turnover after thermal stress. Nevertheless, NADH activities across all designs remained below the wild type level, highlighting the challenge of preserving NADH compatibility in FMOs, which are naturally tuned for NADPH-dependent catalysis.

**Fig 8.**
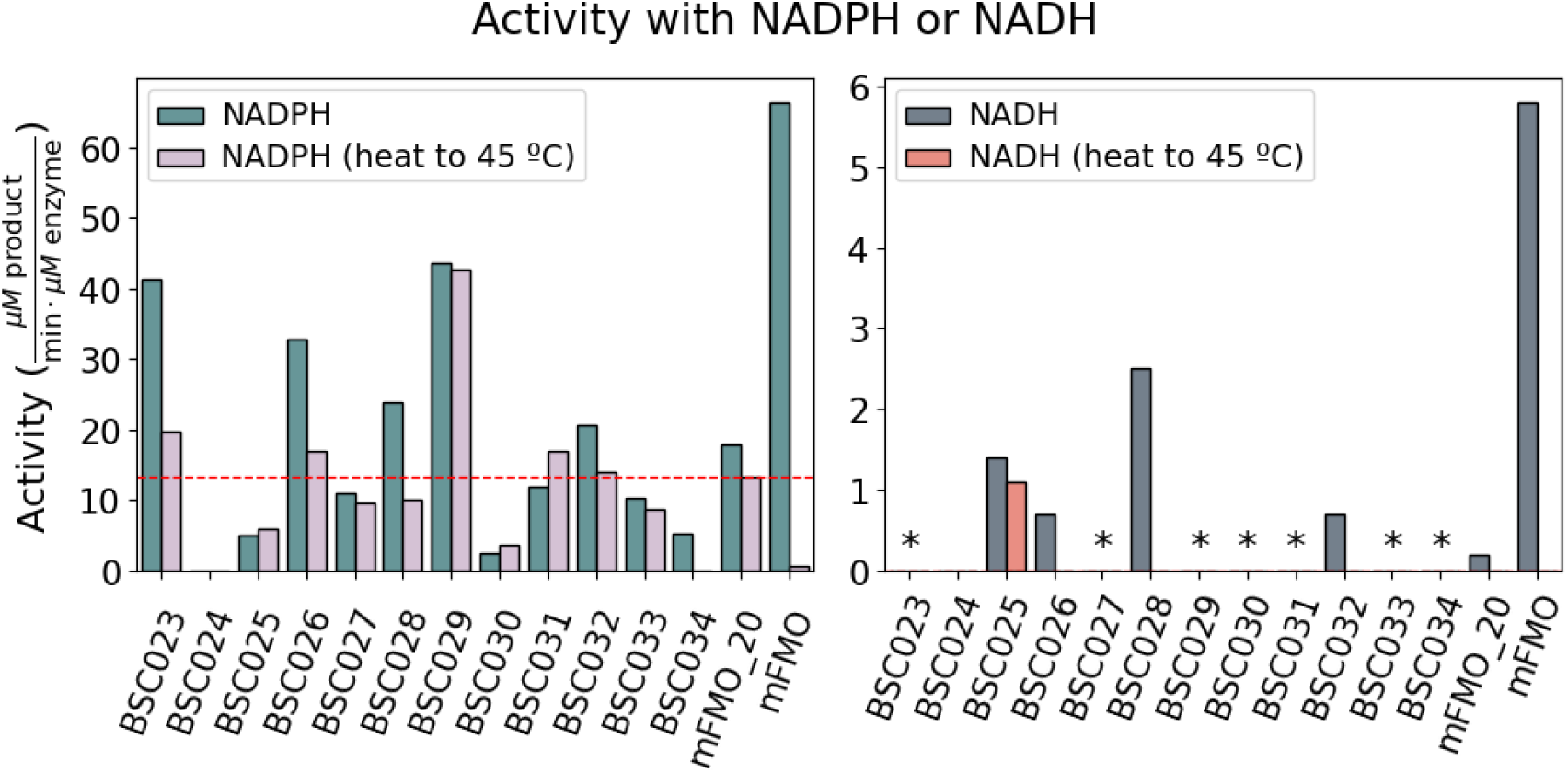
Activity of the mFMO variants obtained from the third optimisation. The indicated mFMO variants (100 nM) were purified, and enzyme activity was calculated using the indicated cofactor (either NADPH or NADH at 100 µM) and TMA (100 µM), monitoring the reaction at 340 nm (n=2) over 1 minute. Variant BSC024 could not be purified and is shown as blank, while asterisks represent purifiable mutants with negligible activity. The red dashed line represents mFMO_20’s activity with NADPH after heating.

Analysis of mutational trajectories clarifies how the algorithm navigated sequence space. The five highest-performing NADPH variants emerged within the first 10 to 20 generations, each accumulating fewer than 16 mutations. In contrast, lower-performing variants appeared in later generations and typically carried more than 19 mutations (S4 Table). Throughout the entire MOO run, fitness increased rapidly during the early generations and plateaued around generation 28 (S6 Fig). This pattern suggests a constrained mutational repertoire, where the most impactful and compatible combinations are sampled early, and subsequent exploration results primarily in incremental changes. Consequently, early generations yielded variants with the greatest retention of NADPH activity and thermal resistance. Later generations, despite accumulating additional mutations, provided minimal further improvement and, in several cases, were associated with reduced activity or stability.

The distribution of mutations across the 12 variants demonstrated that only three substitutions were shared by all designs, while the remaining changes were distributed more heterogeneously. This pattern is consistent with convergence on a small set of stabilising features while preserving sequence diversity. Compared to PROSS, which draws from the same mutant library, the present pipeline introduced a larger set of distinct substitutions (43 unique mutations versus 30 in PROSS; S7 Fig), reflecting broader exploration of the permissible mutational space. Additionally, the selection based on sequence diversity provided a richer pool of candidates for experimental validation while the combination of Potts-guided optimisation with ESM-based ranking offered a more balanced assessment of sequence plausibility than Rosetta energies alone. Collectively, these findings highlight the importance of tuning library composition and integrating evolutionary constraints to minimise the risk of variants that perform well in silico but diverge from natural coevolutionary patterns.

### Molecular Dynamics Analysis of Catalytically Competent Variants

To further probe the mechanistic basis of the experimentally observed differences in NADH activity, we performed microsecond-scale MD simulations of NADH-bound complexes for mFMO, mFMO_20, and two NADH-active designs from the last design campaign (BSC025 and BSC028). For each system, we ran ten independent replicas (1 μs each) and quantified the fraction of frames adopting a hydride-transfer competent geometry, defined by the hydride (H_NADH) to N5 (N5_FAD) distance < 2.7 Å. This reactive-frame occupancy reproduced the experimental ranking of NADH activity before heat treatment (Fig. 9): 7.91% for mFMO, 4.57% for BSC028, 2.24% for BSC025, and 0.14% for mFMO_20 over 10 μs of aggregate sampling.

**Fig 9.**
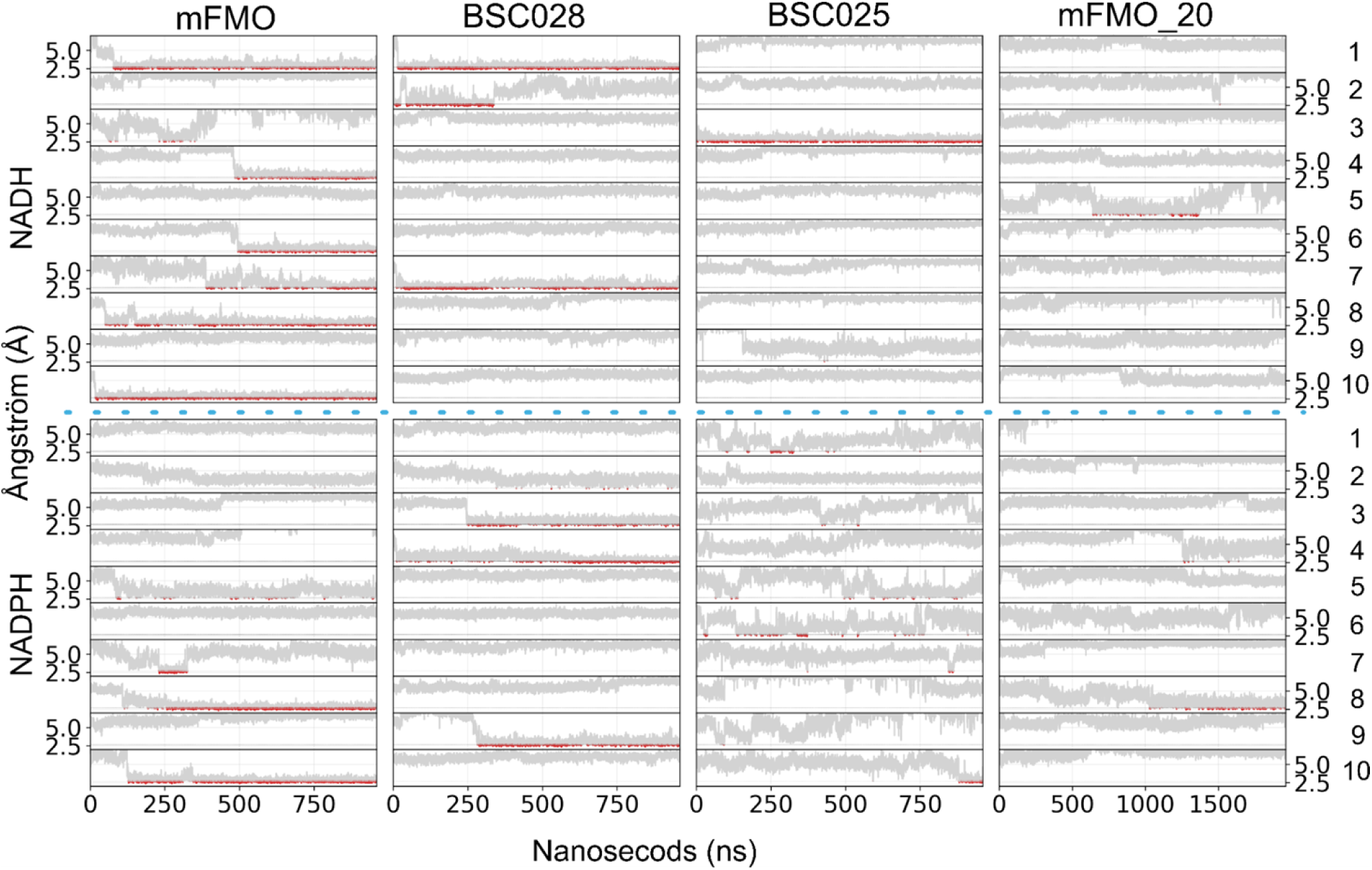
MD simulations of mFMO, mFMO_20, BSC025, and BSC028 in complex with NADH (top four plots) and NADPH (bottom four plots). The x-axis represents simulation time (1000 ns for all systems, and 2000 ns for mFMO_20), while the y-axis shows the distance in Ångström (Å) between the hydride atom of NAD(P)H and the N5 atom of FAD (H_NAD(P)H–N5_FAD). Each system was simulated with 10 independent replicas, displayed as individual line plots and numbered on the right side of the figure. Regions where the catalytic distance falls below the reaction threshold of 2.7 Å are highlighted in red. The blue dotted line separates the NADH from the NADPH simulations.

**Fig 10.**
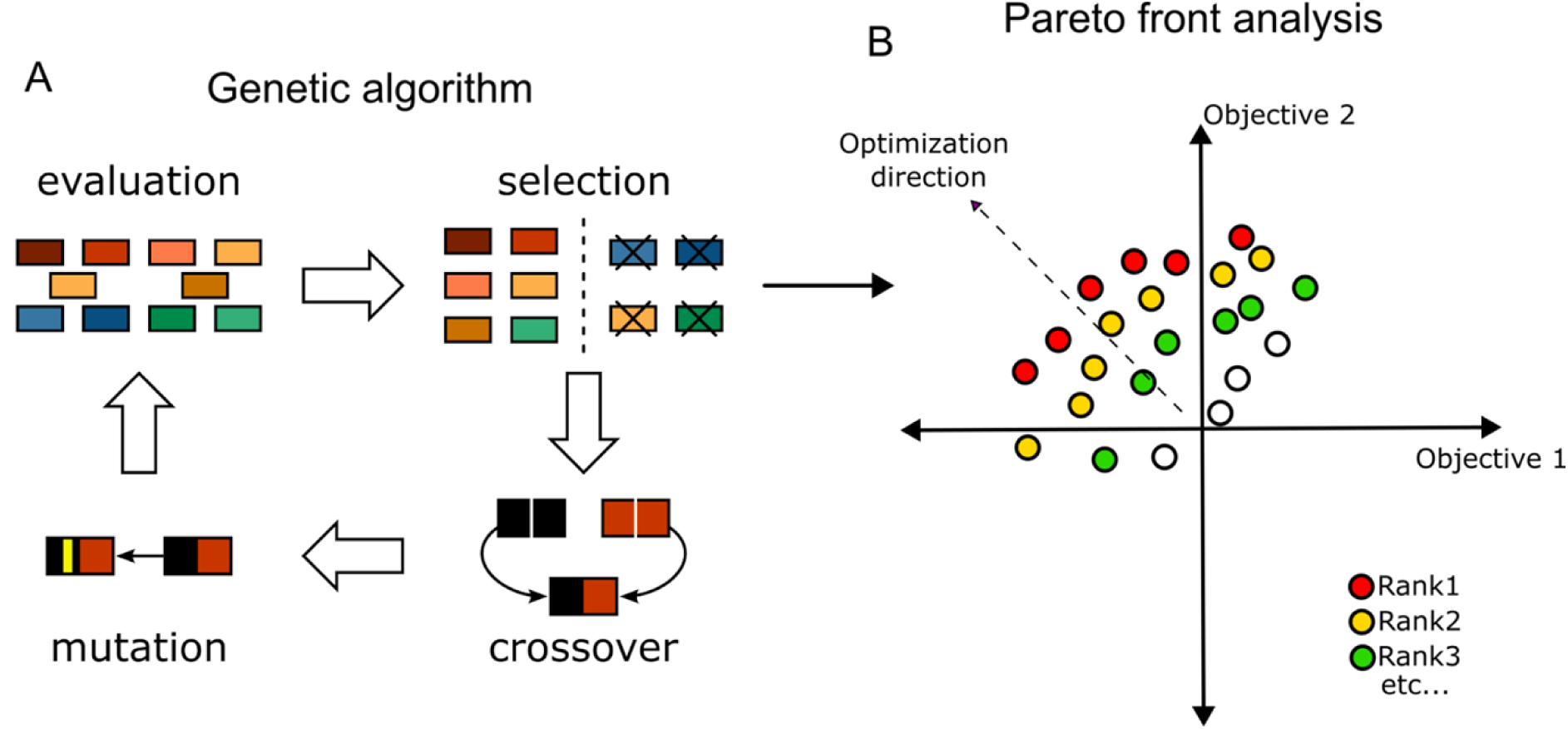
Schematic representation of the multi-objective optimisation algorithm. Panel A illustrates the genetic algorithm’s workflow at each iteration. A population of sequences is first evaluated using multiple fitness functions. A Pareto front analysis is performed to rank the sequences according to their resulting scores. Top-ranked (non-dominated) candidates are selected to serve as parents for the next generation. New sequences are then generated through genetic operators such as mutation and crossover (recombination). Panel B shows the Pareto front ranking process. The first front consists of non-dominated solutions, which are assigned to the highest rank. Subsequent fronts are identified by removing previously ranked sequences and repeating the non-domination analysis until all individuals are ranked.

Except for mFMO, productive geometries were sampled only intermittently: a small subset of replicas accounted for most catalytic-distance events, whereas others remained trapped in non-productive conformations throughout the trajectory. Consistent with this behaviour, the interaction energies associated with frames that satisfy the catalytic-distance criterion were comparable across the NADH-active variants (Fig. S8), indicating that differences arise primarily from the frequency with which reactive geometries are visited, rather than from systematic differences in the stabilisation of the reactive configuration once reached.

We repeated the simulations with NADPH to assess how the same set of enzymes access hydride-transfer competent configurations with their native cofactor (Fig. 9). Using the analogous catalytic-distance criterion (H_{NADPH}-N5_{FAD} < 2.7 Å), the reactive-frame occupancies over 10 μs aggregate sampling were 4.78% for mFMO, 2.91% for BSC028, and 0.79% for BSC025, whereas the PROSS-derived variant, mFMO_20, showed 0% at this timescale. Because mFMO_20 yielded no reactive frames over 10 μs for either cofactor, we extended its sampling to 20 μs; this recovered only rare reactive events (0.14% for NADPH and 0.20% for NADH), indicating that access to catalytic geometries is strongly suppressed and occurs on longer timescales in this variant.

Across variants, experimental NADH activities tracked the MD occupancy of hydride-transfer competent geometries, indicating that access to the catalytic distance window is a primary determinant of turnover. Notably, however, NADPH simulations yielded reactive-frame fractions of the same order as NADH for the same enzymes, whereas experimentally NADPH activities were roughly an order of magnitude higher (see Fig. 8). This discrepancy points to additional cofactor-dependent factors beyond catalytic-geometry accessibility and is likely related to the orientational hypothesis observed during the Monte Carlo simulations (see above). mFMO_20 represented an extreme case: reactive frames were essentially absent at 10 μs for both cofactors and remained rare upon extended sampling (20 μs), yet measurable NADPH turnover was still observed experimentally.

## Discussion

The thermostability of mFMO_20 achieved with PROSS is attributed to interhelical salt-bridge networks spanning helices α6 to α8, which rigidify the scaffold and align with the observed reduction in B-factors [5]. This increased rigidity supports residual NADPH turnover but significantly impairs NADH activity. Our simulations indicate that in mFMO_20, NADH tends to adopt a flipped, ADE-oriented configuration, positioning the NTA away from FAD and preventing hydride transfer. This partially explains the near-complete loss of NADH activity observed experimentally: in the absence of the 2′-phosphate anchor that stabilizes NADPH, NADH is more sensitive to subtle electrostatic and geometric changes within the pocket and can more significantly stabilize nonproductive binding modes.

This interpretation aligns with broader findings in cofactor-specificity engineering, where specificity often results from the overall architecture of the binding pocket and epistatic interactions, rather than from “canonical” residue swaps alone. In *Stenotrophomonas* FMO, restoring a canonical Arg/Thr pair did not enforce strict NADPH dependence, suggesting that cofactor preference depends on the broader pocket context [25]. In *E. coli* glucose-6-phosphate dehydrogenase, NADPH-recognition determinants show non-additive, including antagonistic, interactions, highlighting epistasis as a constraint on the design [26]. Structural evidence also supports the observation that cofactors can adopt a flipped conformation: a NAD-dependent malate dehydrogenase bound to NADP adopts a flipped binding mode, with adenine occupying the catalytic-center position and NTA shifting into the adenine site [27]. Collectively, these studies indicate that cofactor “flipping” and context-dependent residue coupling are plausible mechanisms by which oxidoreductases can enforce or lose NAD(P)H specificity.

Our results show that cofactor specificity and thermostability are linked by at least two distinct constraints. First, at the bound ensemble level, the absence of the NADPH 2′-phosphate increases orientational freedom. Monte Carlo simulations indicate that NADH can access an additional non-productive conformation, which reduces the fraction of catalytically competent molecules, even in the wild type. Second, at the catalytic preorganisation level, activity depends on the frequency of hydride-transfer-competent geometries. MD results suggest that differences in activity between variants, using the same cofactor, arise primarily from changes in the accessibility of these reactive configurations.

Although productive geometries appear to be sampled more frequently for NADH than for NADPH in our simulations, this trend does not reconcile with the lower NADH activity observed experimentally and, if taken alone, would even hint the opposite. This suggests additional factors are at play and implies that the two constraints identified here interact to shape cofactor preference. The mFMO_20 variant is particularly affected by both constraints: reactive geometries are rarely sampled for either cofactor, and the extra orientational freedom of NADH further reduces the productive fraction. We believe that their combined impact likely accounts for the near absence of NADH activity and the presence of NADPH turnover.

To investigate this hypothesis, we first focused on cofactor orientation and sought to reshape NADH binding within the mFMO_20 scaffold. Our MOO selected sequences predicted to stabilize conformations in which the NTA moiety is positioned close to the FAD and disfavour the flipped orientation. While some designs restored measurable NADH turnover, activity remained below wild type levels and NADPH turnover was largely lost. These results indicate that enforcing a preferred binding orientation alone is insufficient to reprogram cofactor use in this scaffold without affecting other requirements for sustained catalysis. These requirements could also be linked to other factors such as electrostatic balance, local dynamics, and steps involved in cofactor recycling, so solutions that address orientation may still disrupt the overall catalytic cycle.

The second design approach focused on improving thermostability from the wild type scaffold while maintaining NADH compatibility by favouring mainly conserved hydrophobic substitutions [28]. Despite favourable Rosetta energy scores of our mutants, this did not result in improved thermal resistance compared to mFMO_20. This discrepancy highlights known limitations of the scoring function. Rosetta’s total score focuses on the folded state, emphasising intramolecular contacts, but does not explicitly model the unfolded or near-unfolded ensembles that often determine thermal stability [20].

In large-throughput studies on de novo αββα proteins, experimental stability correlated with features such as effective core formation and well-defined polar interaction networks. In contrast, Rosetta total energy alone did not reliably predict stabilisation. Critical determinants like helix capping, charge balance, and polar networks can be overlooked when only folded-state energies are optimised [29]. Therefore, while Rosetta energies are useful as a biophysical filter, they may misrank variants if key determinants of thermostability are not included in the objective.

We expanded the optimisation framework to include evolutionary and statistical signals alongside Rosetta energies. Potts model energies capture residue co-variation and encode aspects of epistasis, while ESM language-model likelihoods offer an independent measure of mutational plausibility. Using this composite approach, we achieved measurable gains in thermostability, and one variant retained significant NADH activity, which was not observed with the conservation-restricted strategy. This result represents a better negotiated trade-off rather than a general solution. The predominantly negative outcomes in both approaches suggest that achieving NADH-compatible, heat-resistant function remains challenging due to strict structural and dynamical constraints.

These results help clarify which factors require optimisation. Deep mutational scanning of a FAD-dependent oxidoreductase shows that stabilizing mutations often reduce activity, highlighting a common activity-stability trade-off [16]. Our findings align with this trend and support using evolutionary and statistical metrics to prioritize substitutions that are better tolerated within co-evolved couplings. In simpler oxidoreductases, specificity shifts have sometimes depended on a single position engaging the NADPH 2′-phosphate (e.g., L179S in NADH oxidase; D222Q in formate dehydrogenase), followed by compensatory substitutions to restore activity and stability [30, 31]. In contrast, the present system requires both thermostability and cofactor compatibility in the same scaffold, so multi-signal, context-aware selection is more effective than single-site heuristics.

The added value of Potts models and language models is supported by broader benchmarking. Potts energies capture residue-residue couplings beyond simple conservation [22]. In kinases, Potts correlations with free-energy perturbation and experimental thermostability are comparable to physics-based methods and outperform MM/GBSA, including for nonconservative substitutions [32]. Protein language models (pLMs) such as ESM learn representations linked to biochemical properties and structure, with likelihood scores that correlate with mutational effects across diverse datasets [33]. Zero-shot predictions with ESM-1v perform similarly to family-specific evolutionary models across many datasets, though accuracy decreases in strongly epistatic or multi-mutant contexts [34]. These complementary strengths support a hybrid approach: pLMs offer single-site plausibility priors, Potts models provide explicit epistasis constraints, and integrating both with physics-based scoring can reduce blind spots present in any single method [35]. The Potts-ESM-Rosetta framework therefore serves as a multi-perspective filter, identifying candidates that are both biophysically plausible and evolutionarily more compatible.

The genetic algorithm’s behavior highlights the importance of these signals. The most effective variants appeared within the first 10-20 generations, while later generations accumulated substitutions with limited benefits. This pattern aligns with rapid convergence to local optima in stochastic weak optimisers and demonstrates the value of Pareto fronts for balancing objectives [36]. Variability between runs and premature convergence emphasize the need to incorporate domain knowledge. Integrating Rosetta with Potts and language-model metrics directed exploration toward sequence space that aligns with both biophysical and evolutionary constraints, reducing the risks of purely stochastic methods. These findings suggest a practical approach: use rapid, multi-signal triage to identify Pareto-improving candidates early and avoid late mutation accumulation unless specific mechanistic hypotheses justify it.

Finally, the industrial perspective underscores why NADH compatibility remains an important target. The economic and stability gap between NADPH and NADH is a persistent bottleneck for large-scale applications, and our results highlight the difficulty of achieving this shift through enzyme engineering alone when stability and specificity are tightly coupled. Complementary strategies, therefore, target the cofactor itself. A chemically stabilised analogue, carba-NADP⁺, exhibited a half-life more than 30-fold higher than NADP⁺ at 50 °C while maintaining broad enzyme acceptance [37]. Achieving industrially viable oxygenase platforms will likely require combined approaches for enzyme engineering to broaden cofactor compatibility and cofactor/analogue engineering to improve stability and cost, explicitly recognising that successful solutions must satisfy coupled structural, energetic, and dynamical constraints rather than optimising stability and cofactor preference in isolation.

## Conclusions

This study shows that thermostability and cofactor specificity in this flavin-dependent oxygenase are tightly coupled design constraints. Stabilisation strategies either impaired NADH activity or failed to maintain thermostability under heat stress. A multi-objective framework integrating evolutionary data, protein language models, and physics-based methods improved the identification of stabilising mutations; however, achieving thermostable variants compatible with NADH remains challenging, with only a single design retaining measurable activity. Molecular simulations further showed that this trade-off is shaped by changes in cofactor-binding orientations and in the accessibility of catalytically competent states, helping explain why static structural features alone were insufficient to capture the observed specificity.

Overall, the results suggest that specificity arises from coupled effects between cofactor geometry and global scaffold dynamics, highlighting the need for design strategies that consider cofactor orientations and productive-state accessibility rather than static structural features. Promising directions include explicitly designing and scoring polar or charged interaction networks linked to thermal robustness, incorporating objectives tied to productive-state occupancy or persistence, and pairing enzyme engineering with stabilised cofactor analogues to address cost and thermal degradation. Together, these approaches provide a practical framework for developing oxygenases that balance thermostability with compatibility for cost-effective cofactors.

## Methodologies

### Computational methods

#### The basis for MOO

The genetic algorithm or MOO typically involves three main steps (Fig. 9). The first step is to generate a population of candidate sequences, which can be done using genetic operators such as mutation and recombination. To define a reasonable optimisation space for these operations, a mutation library must be defined, for example, using a Position-Specific Scoring Matrix (PSSM) to determine possible substitutions.

Our program generates a population of 100 sequences by default. Half of these are created by randomly mutating each position with a low probability (0.005) with a residue randomly chosen from the mutant library, while the other half are generated through recombination. During uniform recombination, the algorithm generates a new "child" sequence by randomly selecting, at each position, the corresponding amino acid from one of the parent sequences. Once the initial population is established, the sequences undergo a fitness evaluation, which is the core strength of this algorithm. Multiple fitness functions can be applied, such as Rosetta total energies [20] (similar to PROSS [6]), Rosetta binding energies to ligands or cofactors like NADH, or Potts Energies [22].

To balance potentially conflicting objectives, sequences are ranked using Pareto front analysis to identify the set of non-dominated solutions [38], which are assigned rank 1. After removing these, the next set of non-dominated sequences is assigned rank 2, and so on. From this ranking, a predefined number of sequences (e.g., 50) are selected as parents for the next iteration of the genetic algorithm, prioritising lower-ranked sequences. Importantly, sequences from the previous iteration are also considered, allowing previously discarded sequences to re-enter the population. So in the 50 sequences, there might be sequences of the prior iteration. This framework is similar to the Non-dominated Sorting Genetic Algorithm II (NSGA-II) [39].

In the next iteration, the population is regrown from 50 to the user-defined size of 100 through recombination and mutation. This cycle repeats until the specified number of iterations or generations have been reached.

#### PELE (Protein Energy Landscape Exploration) Simulations

Protein Energy Landscape Exploration (PELE) is a Monte Carlo-based sampling method used to investigate protein-ligand interactions [40, 41]. At each simulation step, the ligand (NADH or NADPH) undergoes small random translations and rotations to explore the conformational space. Protein flexibility is accounted for through normal mode analysis based on the Anisotropic Network Model (ANM). Next, to prevent steric clashes, side chains of residues near the ligand are optimised using a rotamer library. Each proposed conformation is then subjected to truncated Newton minimization and evaluated using the Metropolis acceptance criterion.

The structures of mFMO and mFMO_20 with NADH or NADPH were generated using the AlphaFold 3 server [42] in the NTA conformation, as it can generate more accurate structures than docking tools, whereas the ADE conformation was obtained through docking with Glide [43]. All resulting structures were subsequently prepared using Protein Preparation Wizard at pH 7 [44]. The simulations comprised 50 epochs, with 112 trajectories each. Analysis focused on enzyme-substrate interaction energies and the distance between the C4N atom (the carbon with the hydride) of the nicotinamide ring and the N5 of the FAD in the active conformation, and for the flipped conformation, the C4A atom of the ADE ring and the N5 of FAD. These parameters were automatically computed at every step of the simulation.

#### AsiteDesign

AsiteDesign [21] [https://github.com/masoudk/AsiteDesign] is a Monte Carlo-based protocol (implemented with pyRosetta [45] [DOI: 10.1093/bioinformatics/btq007]) for structure-based engineering of enzymes and their active sites. The Monte Carlo sampling stage involves sampling both the mutagenesis of the defined positions and the ligand conformation, allowing the evolution of the enzyme’s active site toward the substrate orientation. AsiteDesign incorporates enhanced, modular sampling of the substrate and its rotatable bonds, enabling the identification of mutants that might not be found otherwise.

#### Free energy surface analysis of PELE simulation

Free energy surfaces were constructed from PELE simulations using two intermolecular distances measured for each snapshot: FAD-ADE and FAD-NTA. These distances were discretized into a 100 × 100 grid. For each bin, the probability (P) was estimated as:

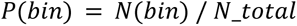

Where N(bin) is the number of snapshots in each bin and N_total is the total number of snapshots. The free energy (F) was then calculated as:

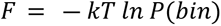

where k is the Boltzmann constant and T the temperature. kT was set to 1.

For each enzyme-cofactor pair, we also analysed the relationship between binding energy and relative total energy. The relative total energy was calculated by subtracting the minimum total energy from the total energy of each pose (Fig. 2). Poses were classified based on their geometry as follows: NTA oriented when the C4N-N5 distance was < 5 Å and ADE oriented when the C4A-N5 distance was < 5 Å.

### MD simulations

#### Simulation set for the selection of mutants in the second design round

Molecular dynamics (MD) simulations were performed for 250 enzyme variants at 60 °C (333.5 K), using three replicas per variant and a simulation length of 150 nanoseconds, all conducted with GROMACS [46] version 2023.3 and the AMBER [47] force field. Structural models were extracted from Rosetta silent files and preprocessed to enable high-throughput thermostability triage: we removed chain B and the NADH cofactor, leaving a single catalytic subunit (chain A) bound to FAD. This simplification was adopted to make screening across 250 variants computationally tractable and to focus the stability metrics (RMSD/RMSF/native contacts) on the fold integrity of the catalytic subunit.

System setup was fully automated using the prepare_proteins library available at https://github.com/Martin-Floor/prepare_proteins.git, specifically the setupMDSimulations function. This pipeline includes system preparation, energy minimization (10 ps), and equilibration steps in the NVT (2 ns) and NPT (0.2 ns) ensembles, followed by a 150 ns production run.

Post-simulation analysis involved averaging the root-mean-square deviation (RMSD), root-mean-square fluctuation (RMSF), and the number of native contacts (Q1) across the three replicas for each sequence and comparing these across variants.

#### Simulation setup for the analysis of the catalytically competent variants

Structures of the mFMO, mFMO_20, BSC025, and BSC028 complexes bound to FAD and NADH were generated with the AlphaFold3 [42] webserver and were used as starting coordinates for molecular dynamics simulations in OpenMM. Each replica was energy-minimized, equilibrated for 0.1 ns in the NVT ensemble while ramping the temperature from 5 K to 298 K, and then equilibrated for 0.2 ns in the NPT ensemble at 298 K and 1 bar. During equilibration, harmonic positional restraints were applied to heavy atoms (excluding solvent and ions) with an initial force constant of 100 kcal mol⁻¹ Å⁻² that was smoothly reduced to zero during NPT. Production simulations were then run unrestrained at 298 K for 1 μs per replica, with 10 independent replicas per system, each using a distinct random seed.

Dynamics were propagated with a Langevin middle integrator using a 2 fs timestep and a 1 ps⁻¹ collision rate, with bonds to hydrogen constrained and rigid water. Periodic box vectors were applied, and long-range electrostatics were treated with PME using a 1.0 nm real-space cutoff and an Ewald error tolerance of 1×10⁻⁴. Pressure was maintained at 1 bar using a Monte Carlo barostat, and coordinates were saved every 100 ps.

Trajectory analysis focused on the hydride-transfer geometry used throughout the manuscript. For each trajectory frame, we measured the distance between the transferring hydride on the NADH nicotinamide ring and the N5 atom of the FAD isoalloxazine ring, denoted d(H_{NADH}–N5_{FAD}). Frames with d(H_{NADH}–N5_{FAD}) < 2.7 Å were classified as hydride-transfer competent, and the fraction of such frames was used as the occupancy reported in Fig. 9. To test whether activity differences reflected binding-strength differences or population differences, protein-NADH interaction energies were recomputed on all frames using the same force-field description and periodic settings.

#### Protein large language models

Protein large language models are trained using a masked language modelling approach, where a single amino acid is hidden, and the model predicts the most probable identity based on the surrounding sequence. After self-supervised training on millions of sequences, the model learns to assign probabilities to each amino acid at every position in a given sequence.

A few studies [34, 48, 49] have demonstrated that protein language models, particularly those from the ESM family, can predict mutation effects on variant fitness, primarily in terms of function and, to a lesser extent, stability, using relative log-likelihoods (ΔLL) compared to the wild type.

Here, we used both ΔLL and ΔP, the sum of probabilities over the sequence relative to the wild type, where higher values imply better mutants.

To compute log-likelihoods (LL) and summed probabilities (P) for each variant, we performed a single forward pass using the ESM-3B model [23], called the wild type marginals [34]. The output logits were transformed into probabilities via a softmax function, ensuring that each row represented a vector of probabilities for all amino acids at that position, with the probabilities summing to 1. The final ΔLL and ΔP scores for a sequence were obtained by summing the log probabilities or probabilities across all positions minus the wild type.

ΔLL is more sensitive to deleterious mutations, as they contribute more significantly to the final score than beneficial ones. Meanwhile, ΔP places greater weight on positive mutations since the deleterious ones have little impact on the final score. They might complement each other when the variants are highly mutated, like in this case, where sequences with up to 30 substitutions were observed.

#### Potts Models

Potts models are probabilistic frameworks derived from multiple sequence alignments, designed to identify co-evolving residues within protein sequences. They achieve this by capturing both site-specific and pairwise frequency patterns observed during training. These models can assign a statistical energy or score to each sequence, where lower Potts energies indicate sequences that better align with the observed evolutionary patterns.

Experimental studies have shown that changes in Potts model scores correlate linearly with changes in folding free energy upon mutation (ΔΔG). Additionally, these scores have been linked to various experimental measures of protein fitness, including thermostability, enzyme activity, and overall stability [22].

#### Rosetta

Rosetta’s modelling suite is central to the program. All energies were calculated using the scoring function ref2015. The ExposedHydrophobics filter and the SaltBridgeFeatures were used to identify exposed hydrophobic atoms (which reflect hydrophobic packing) and to count salt bridges, respectively. These were used because they are generally considered relevant indicators of thermostability [24].

### Experimental Methods

#### Enzyme expression and purification

Genes encoding the novel mFMO variants were purchased from Twist Bioscience (San Francisco, USA) as fragments with flankers containing SapI restriction sites and FX cloned into a pBAD plasmid, essentially as described here [3]. This plasmid is under the control of an arabinose-inducible promoter and encodes a C-terminal 6xHis-tag. MC1061 chemically competent *E. coli* were transformed with the plasmids, and the cells were grown to an OD of 0.6 at 37 °C. They were induced by adding 0.1% arabinose and expressed overnight at 20 °C. After expression, bacteria were harvested by centrifugation and stored at -20°C until purification. The enzymes were purified as described in [5]. Briefly, bacterial cells were lysed using freeze/thaw cycles and sonication; the lysate was cleared by centrifugation, then loaded onto a nickel-NTA column and washed with low imidazole before being eluted with high imidazole. The enzymes were desalted on a PD-10 column and upconcentrated using a molecular weight cutoff filter (30 kDa). After upconcentration, the enzymes were diluted to 10% glycerol and aliquoted. The aliquots were frozen at -80 °C and then stored at -20 °C. Enzyme concentration was measured by the 660 nm protein assay (Pierce). mFMO and mFMO_20 were expressed and purified in parallel with the novel variants.

#### Enzyme assays and heat treatment

Heat treatment was performed by diluting aliquoted enzymes to 1 µM in reaction buffer (50 mM tris-HCl, pH 8.0 and 50 mM NaCl) to a final volume of 250 µl, then incubating in a water bath at 45 °C for 30 minutes. The enzymes were then chilled on ice and immediately used for enzyme assays. Enzymes without heat treatment were simply diluted to 1 µM and kept on ice.

The enzyme assays were conducted essentially as described [5], with some modifications that reduced the preincubation time and the cofactor concentrations. Briefly, 840 µl reaction buffer (50 mM tris-HCl, pH 8.0 and 50 mM NaCl) was mixed with enzyme (final concentration 50 or 100 nM), cofactor (either NADH or NADPH, final concentration 100 µM), and incubated for 00:30 seconds. Then, the reaction was started by adding 100 µl TMA (1 mM, final concentration 100 µM). The mixture was well mixed, and the absorbance at 340 nm was measured immediately in a spectrophotometer for 1 minute. All enzymes showed a linear change in absorbance over this period. Enzymes were tested with both cofactors in duplicates. Concentration changes in NADH/NADPH were calculated using the Beer-Lambert equation, based on the absorbance change over 1 minute and assuming a molar extinction coefficient of 6.22 mM^-1^ cm^-1^ for NADH and NADPH. The activity is reported as µM product formed/min/µM enzyme, or min^-1^.

## Supporting Information

### Supplementary figures

**S1 Fig.**
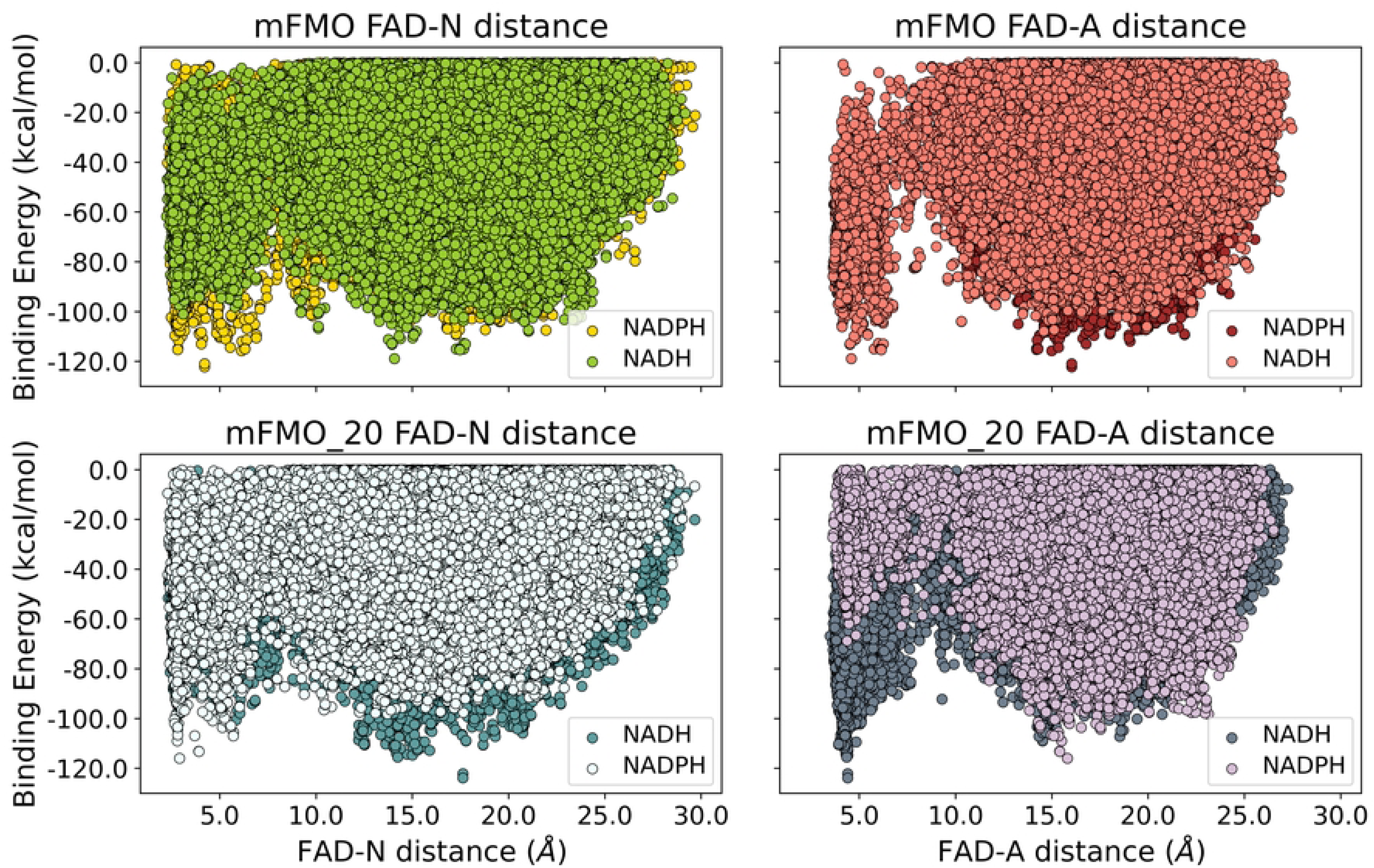
PELE simulations of mFMO and mFMO_20 with NADH or NADPH as cofactors. Panels A and B correspond to mFMO, while C and D show results for mFMO_20. In A and C, the x-axis represents the distance between the nicotinamide ring of the cofactor (NADH and NADPH) and the isoalloxazine ring of FAD. In B and D, the x-axis shows the distance between the adenosine ring of the cofactor and FAD. The y-axis in all plots indicates the binding energy between the enzyme and the cofactors. Panel D reveals that the adenine portion of NADH is more stable inside the active site of mFMO_20 than the nicotinamide part shown in C, as it reaches a lower energy minimum. While A and B show that the two conformations of NADH are equally stable in the FMO’s active site.

**S2 Fig.**
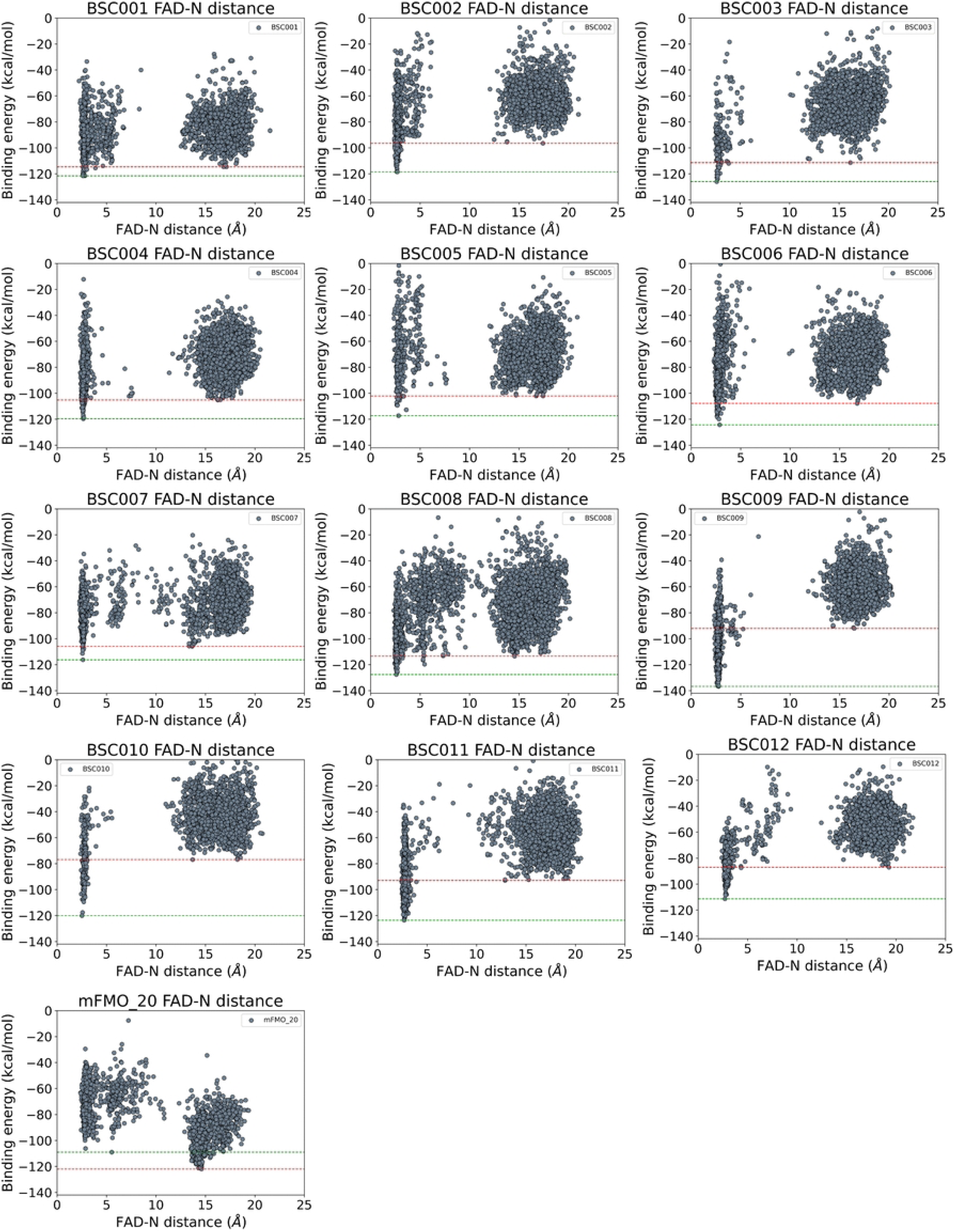
PELE simulations of the variants derived from the first MOO run. The red line indicates the minimum binding energy for conformations with adenosine-oriented NADH binding, while the green line represents the minimum binding energy for nicotinamide-oriented conformations.

**S3 Fig.**
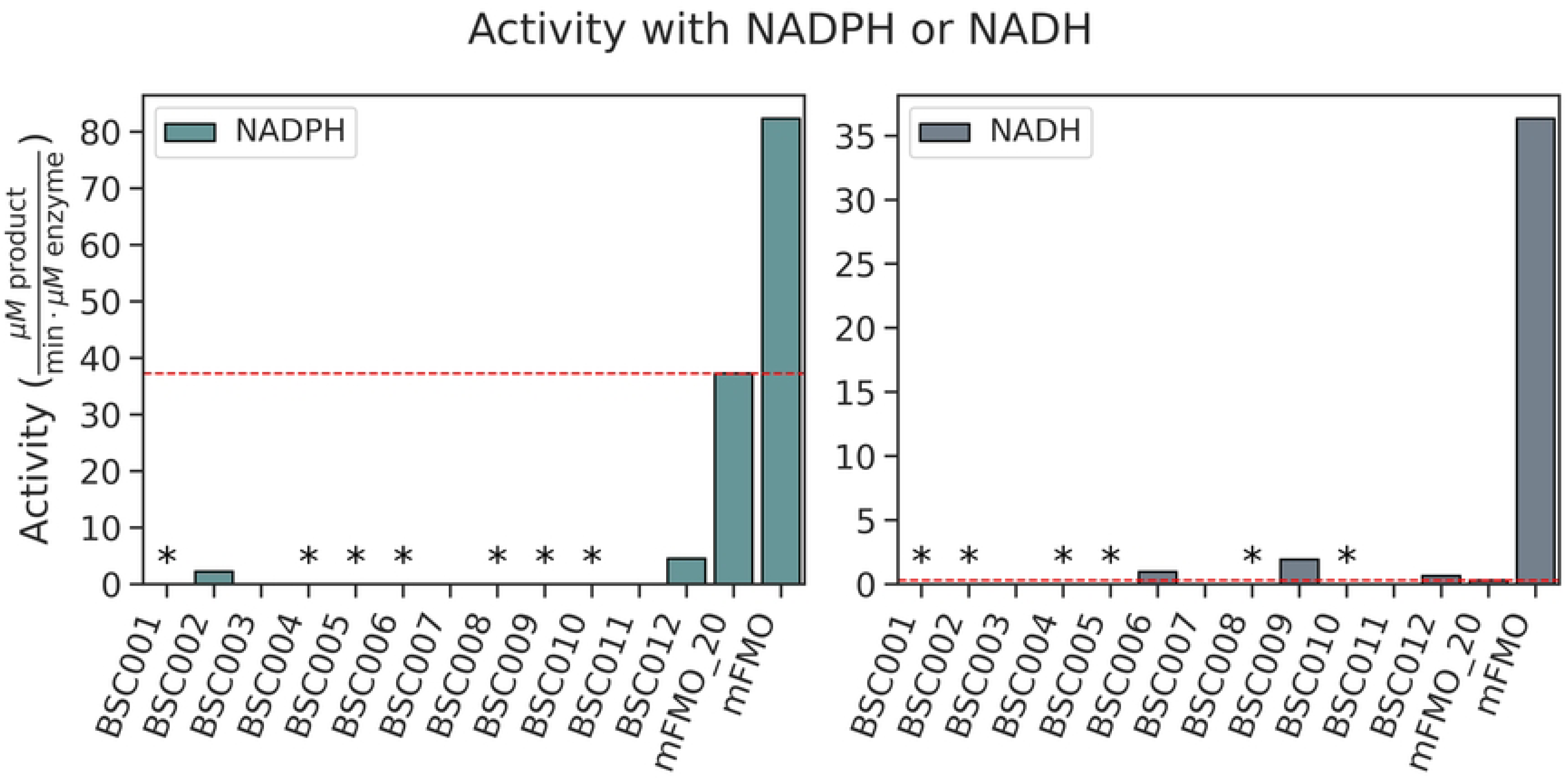
Activity of the variants from the first MOO run. The indicated enzymes (50 nM) were purified, and their velocity was measured using the indicated cofactor (either NADPH or NADH at 100 µM) and TMA (100 µM), monitoring the reaction at 340 nm (n=2) over 2 minutes. BSC001 was designed to target residues around the phosphate-binding site; BSC002-BSC008 were designed using AsiteDesign, and BSC009-BSC012 were designed using multi-objective optimisation. Variants that are not expressed are shown as blanks; asterisks indicate expressed variants with no detectable or negligible activity. The red line represents mFMO_20’s activity with each cofactor.

**S4 Fig.**
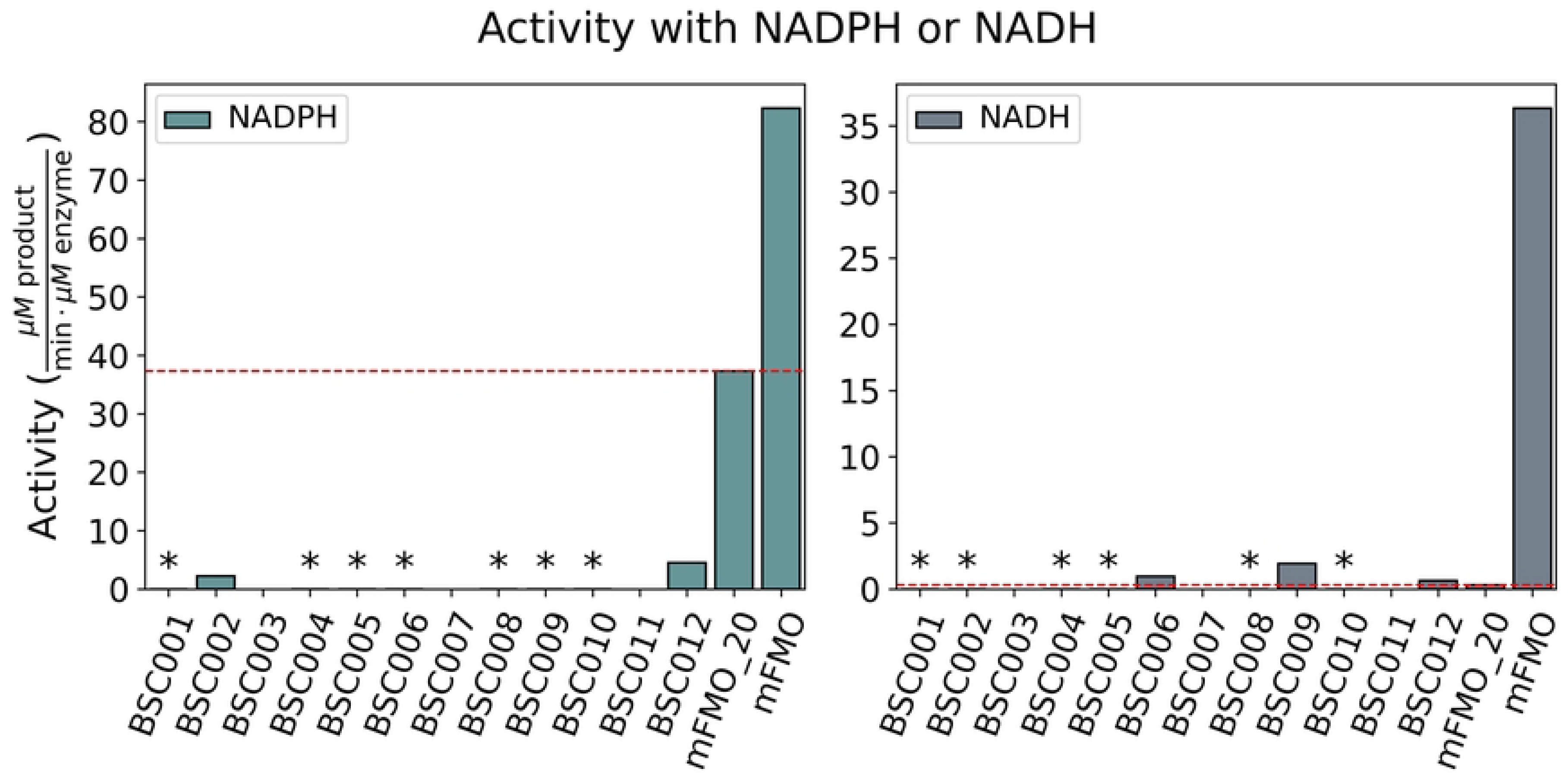
Comparison of Rosetta total energies between BSC mutants and PROSS mutants. The Rosetta total scores of the selected BSC mutants are in blue, while the PROSS mutants are in green. The BSC mutant BSC020 and the PROSS mutant mFMO_20 were shown separately. The Y axis shows the number of mutations, and the X axis shows the delta total energies between the mutants and the wild type mFMO.

**S5 Fig.**
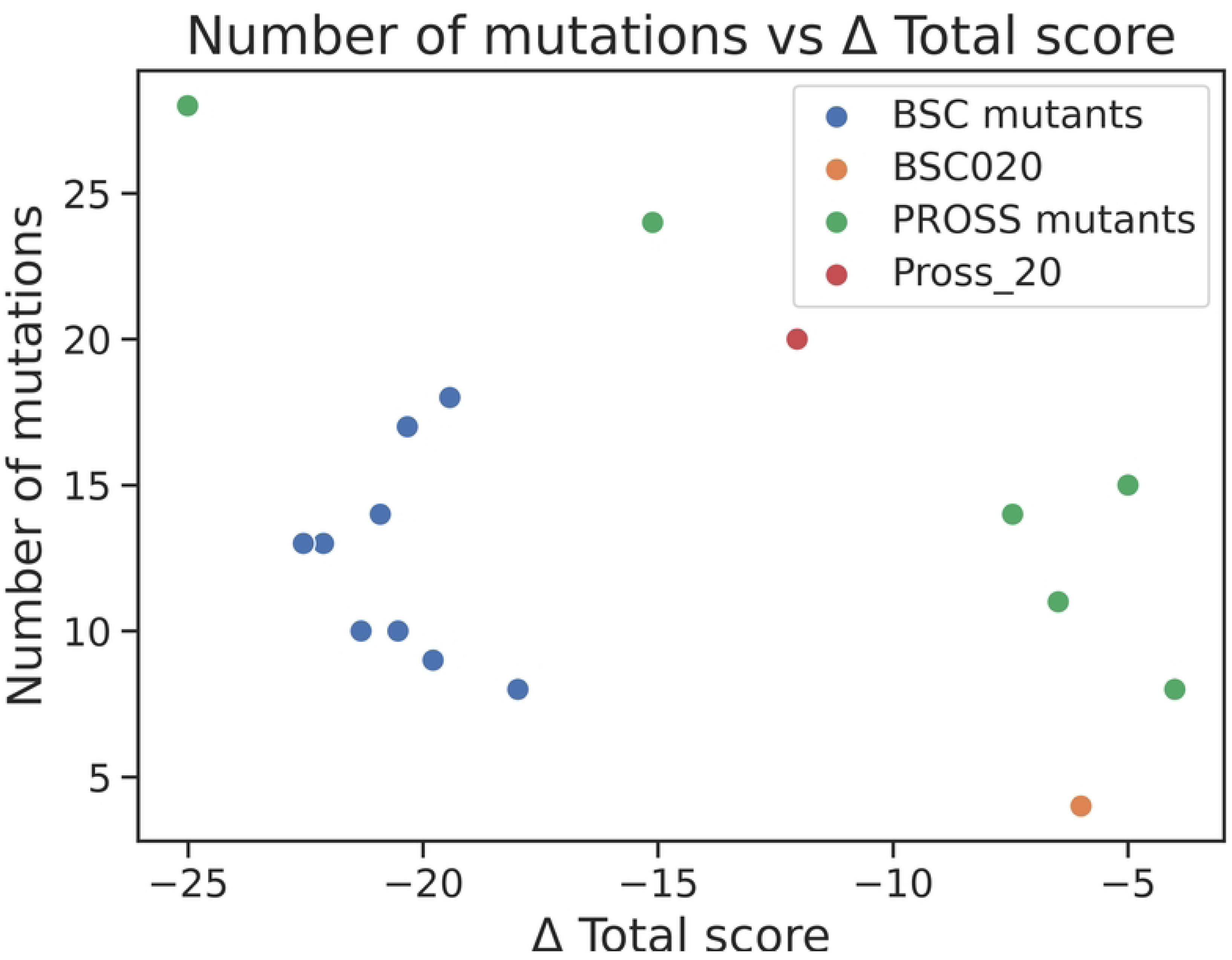
Boxplot of the Rosetta total scores across the last 2 design runs.

**S6 Fig.**
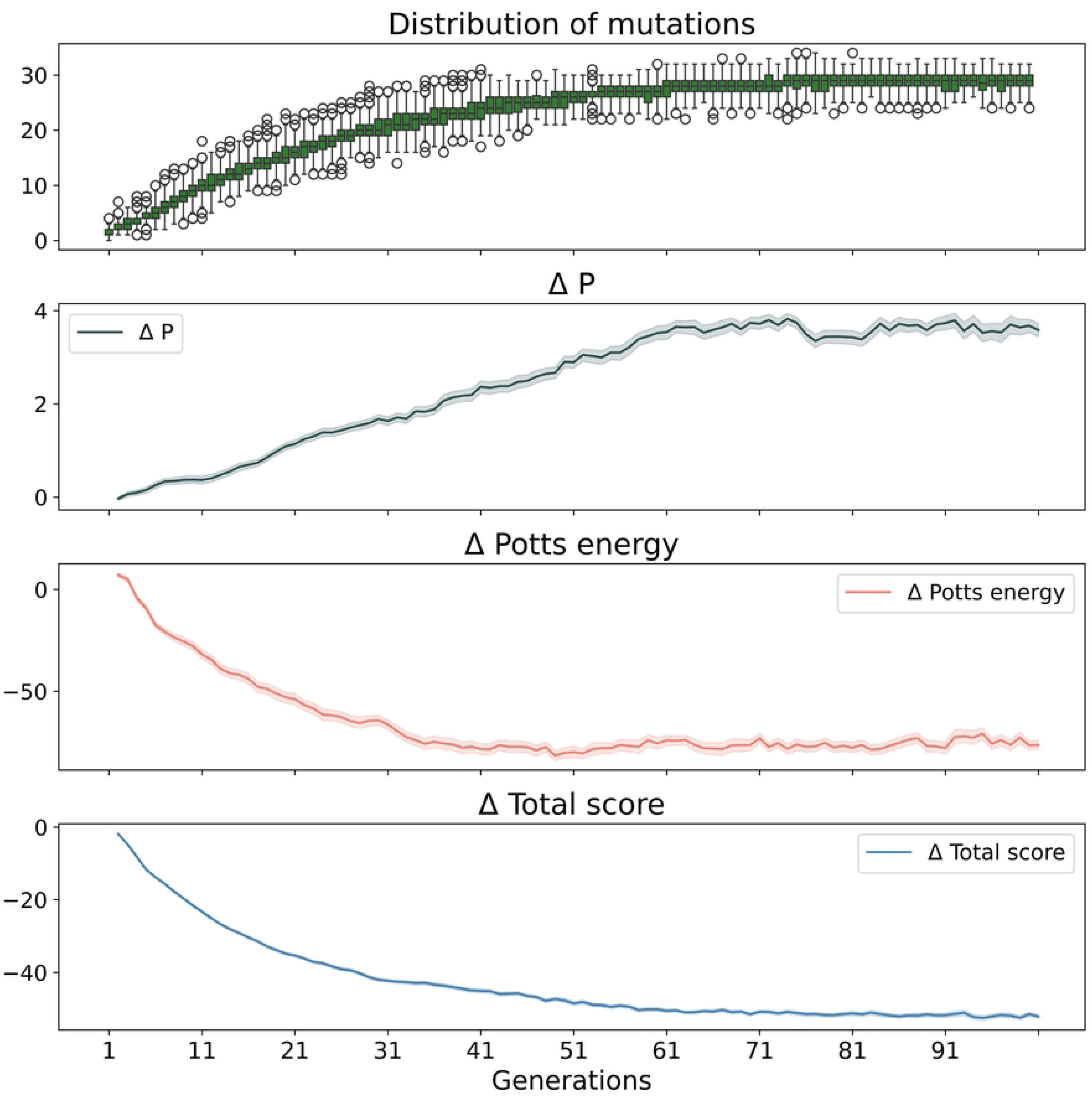
Changes in fitness scores over successive generations of the MOO run. The X-axis represents the number of iterations or generations, while the Y-axis shows the number of mutations and the corresponding scoring metrics: ESM2 probabilities, Potts energies, and Rosetta total energies. It shows steep improvements in early generations, followed by a plateau as the library’s mutational space is exhausted.

**S7 Fig.**
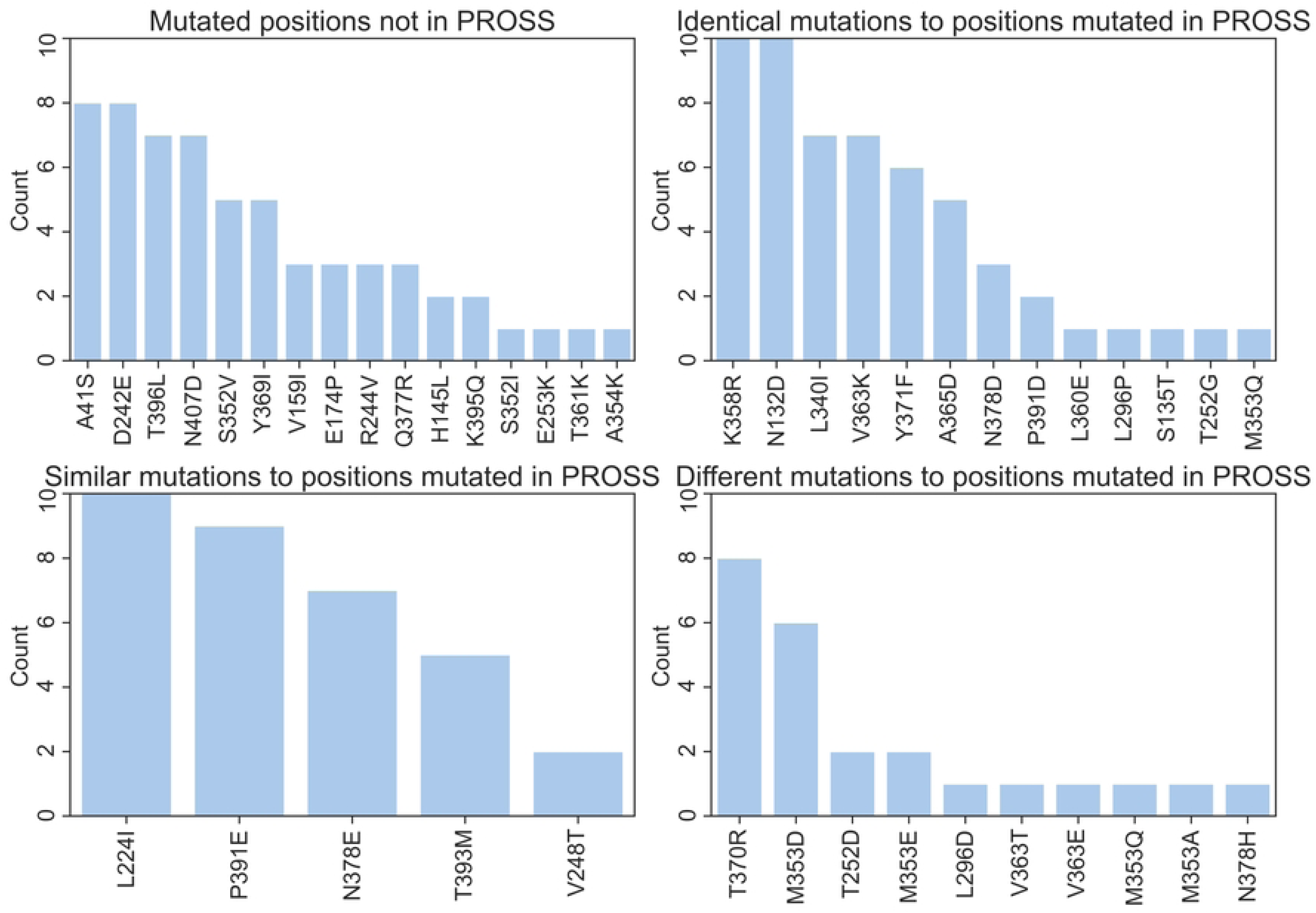
Mutation frequency comparison between selected variants and PROSS mutants. “Mutated positions not in PROSS” shows those positions that are only mutated in our set. “Identical mutations” are those shared with PROSS mutants. “Similar mutations” refer to positions mutated in PROSS as well, but to a residue that is only similar to the PROSS mutant, for example, hydrophobic to another hydrophobic residue. While “different mutations” mean the same position is also mutated in PROSS, but to a completely different residue. The X-axis is the number of sequences with a given mutation, while the Y-axis is the different mutations.

**S8 Fig.**
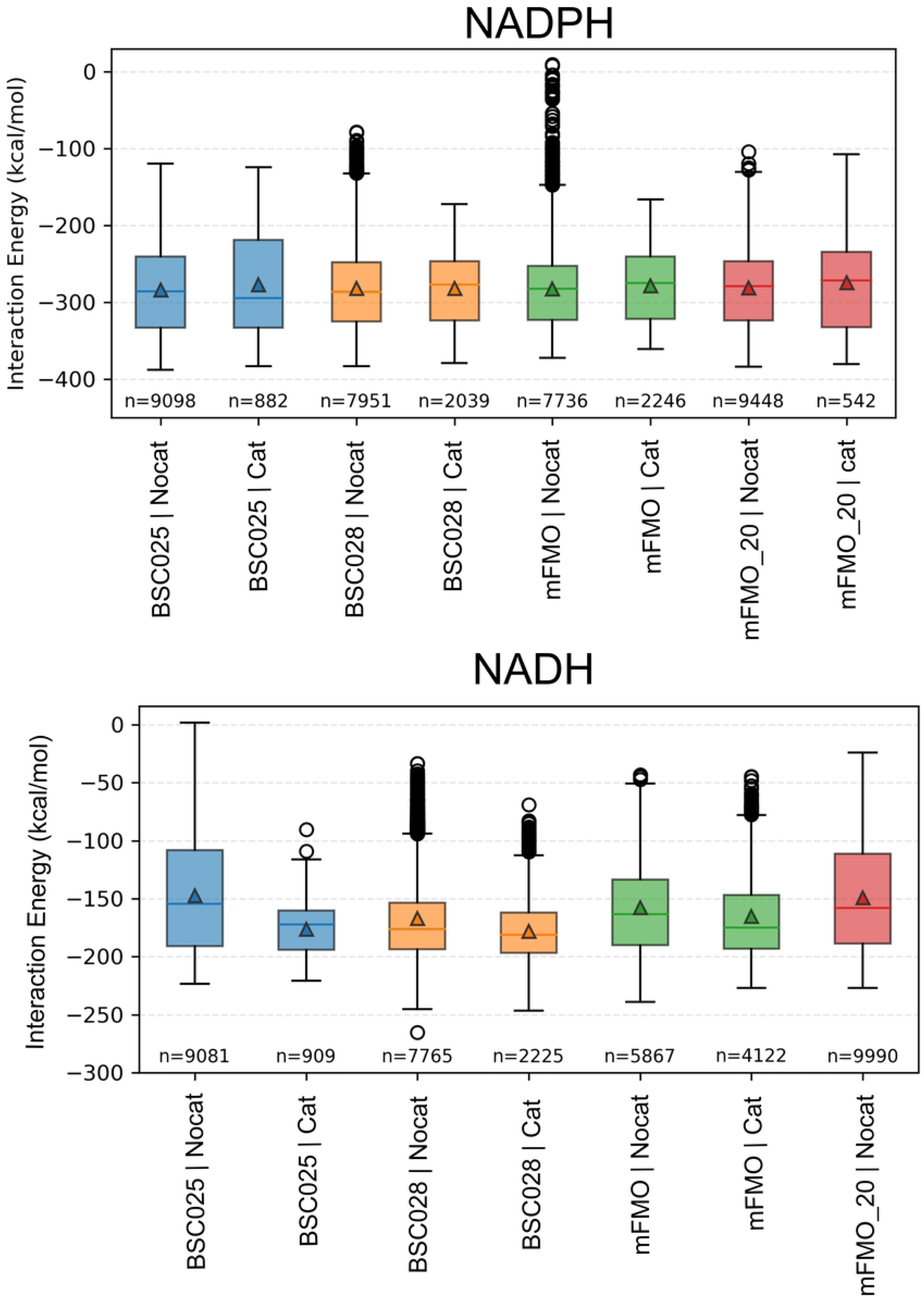
Boxplot of interaction energies obtained from MD simulations of mFMO, mFMO_20, BSC025, and BSC028. For each simulation, conformations in which the distance between the catalytic atoms was within the 2.7 Å threshold were classified as “Cat”, while all remaining conformations were classified as “NoCat”. In the case of mFMO_20, no conformations satisfying the catalytic distance criterion were observed, and therefore, it is not shown. *N* indicates the number of poses in each class. The y-axis represents the interaction energy (kcal/mol), and the x-axis indicates the enzyme variant.

### Supplementary tables

**S1 Table. The activity of mFMO_20 mutants from the first optimisation run is expressed as the turnover number ([µM product formed min-1 µM enzyme-1] or [min-1])**.

**S2 Table. The activity of mFMO mutants from the second optimisation run expressed as velocity ([µM product formed min-1 µM enzyme-1] or [min-1] and the mutated positions**. The indicated mFMO variants’ activity was measured using the indicated cofactor (either NADPH or NADH at 100 µM) and TMA (100 µM), monitoring the reaction at 340 nm (n=2) for 1 minute, either after heating at 45 °C or after no heat treatment. Variants 14, 15, 17, and 21 were not tested after heating.

**S3 Table. The activity of mFMO mutants from the third optimisation run expressed as velocity ([µM product formed min-1 µM enzyme-1] or [min-1] and the mutated positions**. The indicated mFMO variants (100 nM) were purified, and enzyme velocity was measured using the indicated cofactor (either NADPH or NADH at 100 µM) and TMA (100 µM), monitoring the reaction at 340 nm (n=2) over 1 minute. Variant BSC024 could not be purified.

**S4 Table. Generation number and mutation count for the twelve BSC variants.**

### Supplementary methods

**S1 Methods: Analysis of the PELE induced-fit simulations**

**S2 Methods: Description of the mutant design process using AsiteDesign, the first multiobjective optimization (MOO) run, and the targeted approach.**

**S3 Methods: Mutant design process for the second optimisation run.**

**S4 Methods: Selection of mutants for the third optimisation run.**

## Appendix

**S1 Appendix: The mutant library used by PROSS to design mFMO_20**

---

**S4_Method_Fig2.**
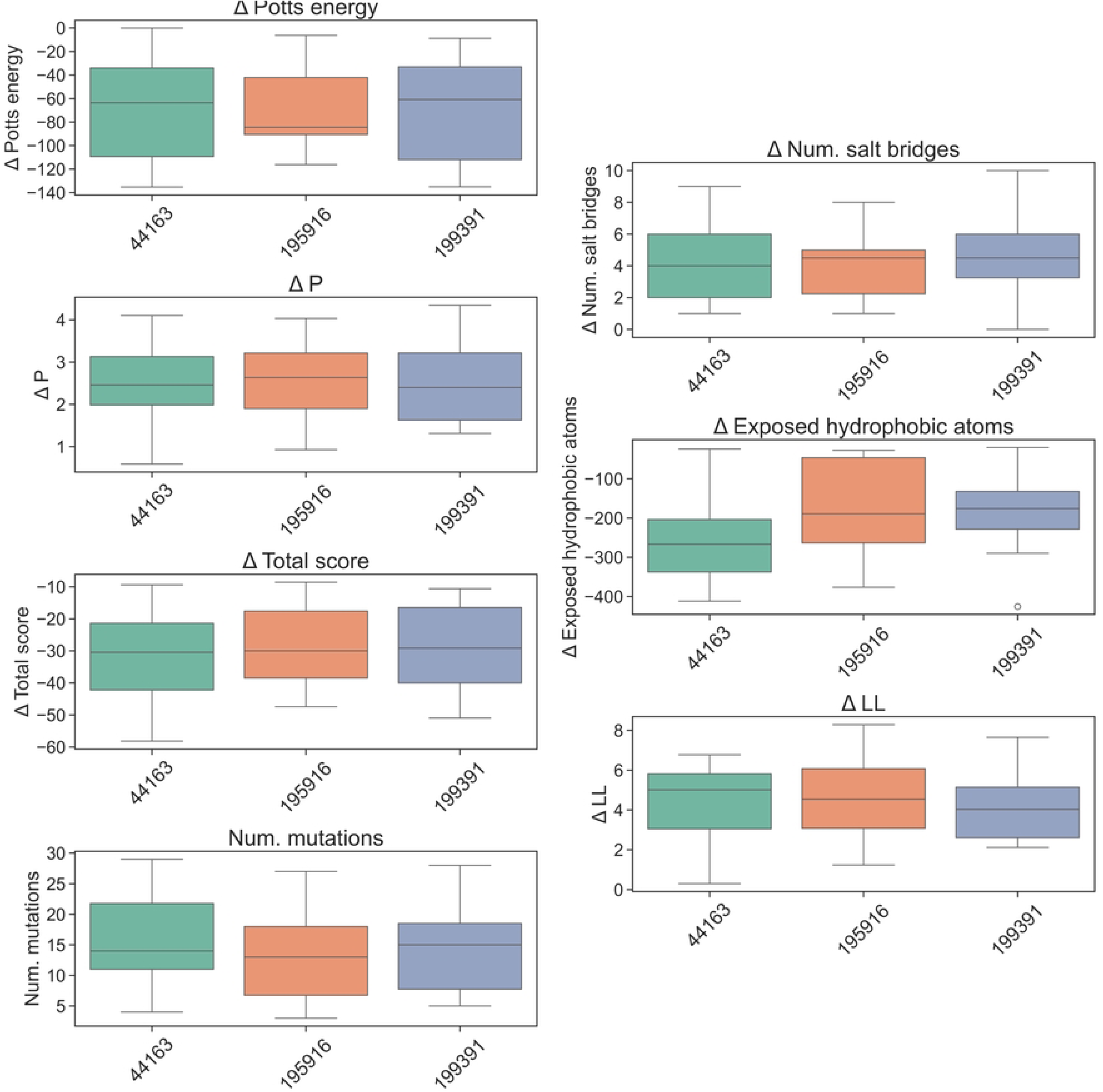

**S4_Method_Fig2.**
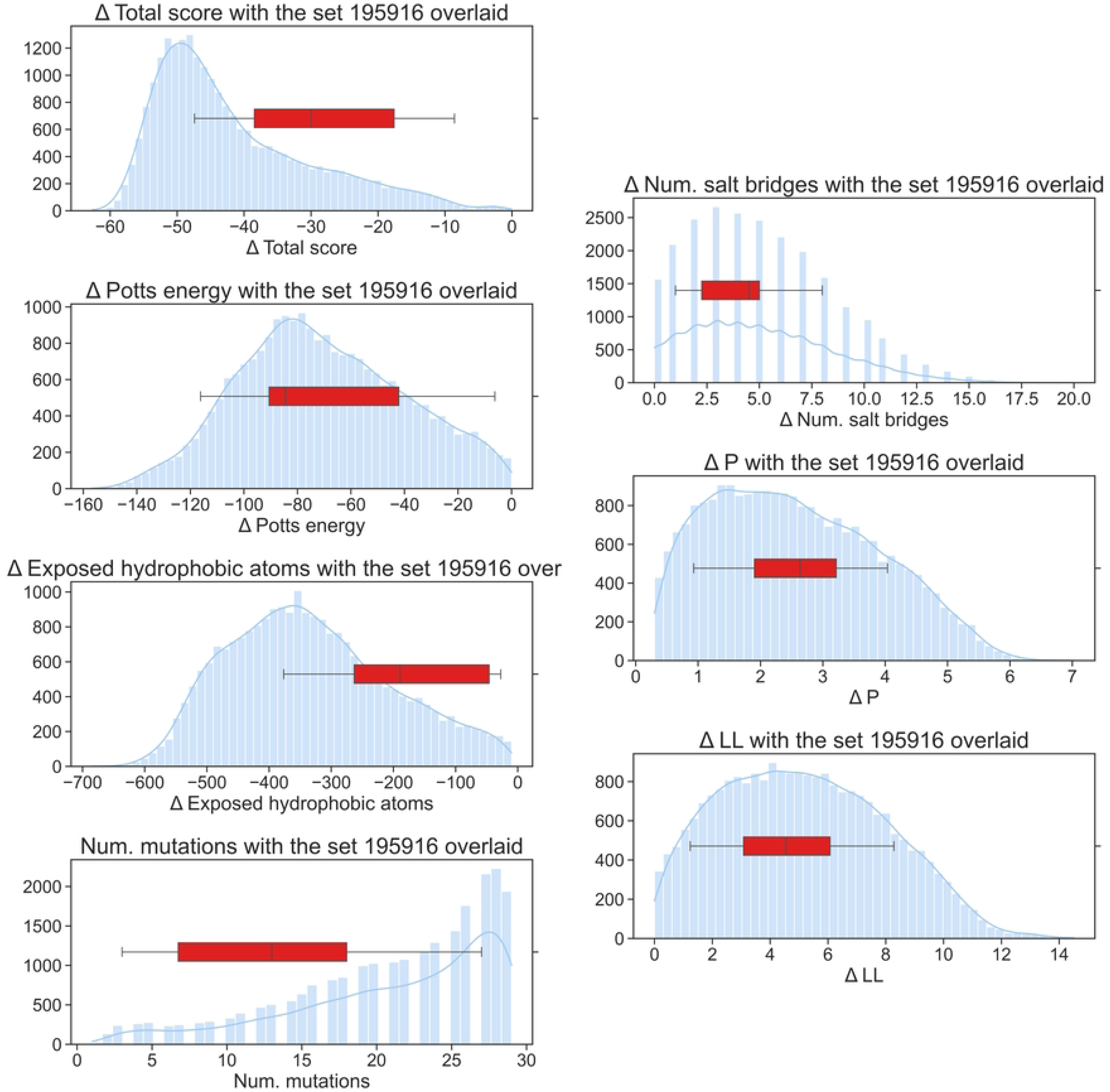

